# The transaminase-ω-amidase pathway is a redox switch in glutamine metabolism that generates α-ketoglutarate

**DOI:** 10.1101/2024.08.28.610061

**Authors:** Niklas Herrle, Pedro F. Malacarne, Timothy Warwick, Alfredo Cabrera-Orefice, Yiheng Chen, Maedeh Gheisari, Souradeep Chatterjee, Matthias S. Leisegang, Tamim Sarakpi, Sarah Wionski, Melina Lopez, Ina Koch, Marcus Keßler, Sabine Klein, Frank Erhard Uschner, Jonel Trebicka, Steffen Brunst, Ewgenij Proschak, Stefan Günther, Mónica Rosas-Lemus, Nina Baumgarten, Stephan Klatt, Thimoteus Speer, Ilka Wittig, Marcel H. Schulz, J. Brent Richards, Ralf Gilsbach, Travis T. Denton, Ingrid Fleming, Luciana Hannibal, Ralf P. Brandes, Flávia Rezende

## Abstract

Oxidative stress is caused by short-lived molecules and metabolic changes belong to the fastest cellular responses. Here we studied how the endothelial cell metabolome reacts to acute oxidative challenges (menadione or H_2_O_2_) to identify redox-sensitive metabolic enzymes. H_2_O_2_ selectively increased α-ketoglutara<u>ma</u>te (αKGM), a largely uncharacterized metabolite produced by glutamine transamination and a yet unrecognized intermediate of endothelial glutamine catabolism. The enzyme nitrilase-like 2 ω-amidase (NIT2) converts αKGM to α-ketoglutarate (αKG). Reversible oxidation of specific cysteine in NIT2 by H_2_O_2_ inhibited its catalytic activity. Furthermore, a variant in the *NIT2* gene that decreases its expression is associated with high plasma αKGM level in humans. Endothelial-specific knockout mice of NIT2 exhibited increased levels of αKGM and impaired angiogenesis. Knockout of NIT2 impaired endothelial cell proliferation and sprouting and induced senescence. In conclusion, we show that the glutamine transaminase-ω-amidase pathway is a metabolic switch in which NIT2 is the redox-sensitive enzyme. The pathway is modulated in humans and functionally important for endothelial glutamine metabolism.

## Introduction

Oxidative stress is a hallmark and a potential driver of cardiovascular diseases. Reactive oxygen species (ROS) are a heterogeneous class of molecules, which differ in their reactivity, biological targets and functional relevance. Nitric oxide, superoxide anions (O_2_^•-^) and hydrogen peroxide (H_2_O_2_) are of particular importance due to their relevance in cellular signaling.^1^ Cellular responses to an acute challenge with ROS have been mainly studied regarding signal transduction and gene expression. Because ROS are short-lived molecules, fast changes in the cellular metabolome are at the forefront response to oxidative stress.^2^ The characterization of redox regulation of metabolism remains incomplete, despite of significant advances in the field of metabolomics. Well established examples of redox-control of metabolism are largely restricted to the best studied metabolic pathways such as the superoxide-sensitive iron-sulfur clusters of aconitase and isocitrate dehydrogenase (ACO1, IDH1, tricarboxylic acid (TCA) cycle)^3^ as well as glucose-6-phosphatase dehydrogenase (G6PD, pentose phosphate pathway),^4^ glyceraldehyde 3-phosphate dehydrogenase (GAPDH, glucose oxidation)^5^ and pyruvate kinase M2 (PKM2, glycolysis).^6^

The metabolic plasticity of endothelial cells is unique; as depending on their environment they can be proliferative, quiescent, stationary or migratory and, thus, are exposed to high or low partial pressure of oxygen. Endothelial cells are also a prime site of inflammation and, as such, are exposed to changes in their redox environment or even overt oxidative stress. Although an oversimplification, it has been reported that endothelial cells re-generate ATP mainly via glycolysis while oxidizing only < 1 % of the pyruvate generated in TCA cycle.^7–9^ In human umbilical vein endothelial cells (HUVEC), the glycolytic flux is more than 200-fold higher than glucose oxidation in the electron transport chain.^10–12^ This is supported by the fact that endothelial cells have few mitochondria and generate nitric oxide, which inhibits mitochondrial respiration. This inhibition reduces oxidative phosphorylation, oxygen consumption, and mitochondrial ROS generation.^13^ Thus, endothelial cells utilize glutamine as an anaplerotic source of carbon for the biosynthesis of nucleic acids, lipids and other building blocks required for proliferation.^14,15^ Within the cell, glutamine is converted to glutamate by glutaminase (GLS1) and thereafter to α-ketoglutarate (αKG) by glutamate dehydrogenase (GLUD1) or by transamination with a suitable α-keto acid substrate. This reaction is known as the glutaminase I pathway and αKG is a central TCA cycle intermediate and at the crossroads of several metabolic processes. Based on the metabolic features of endothelial cells we analyzed the endothelial metabolome in response to an acute oxidative challenge with menadione (to generate O_2_^•-^) or extracellular H_2_O_2_. We observed that H_2_O_2_ selectively unmasked a redox-sensitive and functionally important non-canonical pathway for the generation of αKG from glutamine.

## Results

### H_2_O_2_ increases α-ketoglutara<u>ma</u>te levels in endothelial cells

To assess the metabolic response of endothelial cells to menadione and H_2_O_2_, we utilized our previously published dataset, which compared the time-resolved responses to various types of ROS.^16^ Menadione, a redox cycler that generates intracellular O_2_^•-^, led to a significant decrease in isocitrate levels consistent with the known inhibition of aconitase by superoxide anions.^17^ However, the levels of downstream metabolites of αKG were not affected. This can be explained by the fact that glutamine, through GLS1, replenishes carbons via αKG into the TCA cycle (**Fig. 1A, 1C**). Strikingly, H_2_O_2_ selectively decreased the levels of αKG and its downstream metabolites in the TCA cycle but also conversely increased the levels of α-ketoglutara<u>ma</u>te (αKGM, **Fig. 1B-E**). This largely uncharacterized metabolite is rarely mentioned in the literature because it was not commercially available and, thus, not measured by targeted mass spectrometry-based analysis (LC-MS/MS).^18^ αKGM is formed by transamination of glutamine by the transaminase enzymes KYAT1 and KYAT3 (kynurenine aminotransferases, previously annotated in the human genome as GTK and GTL for glutamine transaminase of kidney and liver, respectively).^19^ αKGM can be converted to αKG by de-amidation and if not metabolized it forms a stable lactam: 2-hydroxy-5-oxo-proline. Since H_2_O_2_ increased αKGM and decreased αKG in endothelial cells, this reaction likely involves an enzyme that is redox-sensitive and inhibited by H_2_O_2_.

**Figure 1:**
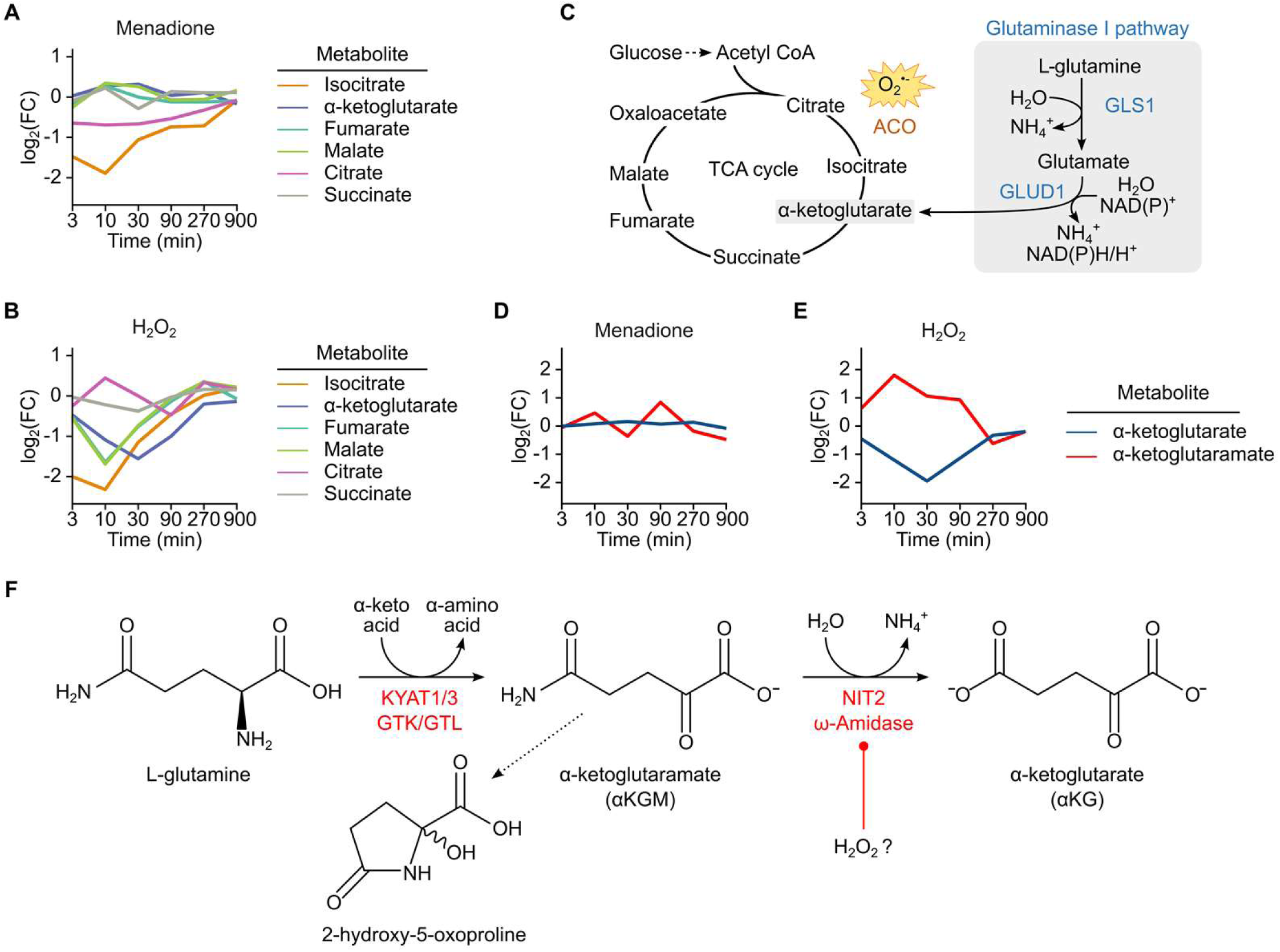
Exposure of endothelial cells to H_2_O_2_ increases α-ketoglutara<u>ma</u>te (αKGM). Mean of log_2_ fold change in TCA cycle metabolites (untargeted metabolomics) of human umbilical endothelial cells (HUVEC) exposed to either menadione (5 µM) (A) or H_2_O_2_ (300 µM) (B). C: Superoxide oxidizes the iron-sulphur cluster in aconitase, however, the glutaminase I pathway can replenish α-ketoglutarate (αKG) into the TCA cycle through the action of glutaminase 1 (GLS1). D-E Changes in αKGM and αKG in HUVEC after exposure to menadione or H_2_O_2_. F: Reactions of the glutamine transaminase-ω-amidase (GTωA) pathway. ACO: Aconitase GLUD1: Glutamate Dehydrogenase 1 KYAT1: Kynurenine aminotransferase 1 or Glutamine transaminase of kidney KYAT3: Kynurenine aminotransferase 3 or Glutamine transaminase of liver NIT2: Nitrilase-like 2, ω-amidase

### De-amidation of αKGM to αKG is catalyzed by NIT2

The de-amidation of αKGM to αKG is catalyzed by the nitrilase-like 2, ω-amidase enzyme, NIT2. The non-canonical reaction cycle to generate αKG from glutamine is referred to as glutamine transaminase-ω-amidase pathway (GTωA) (**Fig. 1F**).^20^ This pathway was proposed in the 1950s^21,22^ but has been only partially characterized.

αKGM is contained in the untargeted metabolite panel of Metabolon^®^ (as determined by its fragmentation pattern) but LC-MS/MS confirmation on the basis of standards has not been performed. We therefore generated αKGM in its pure form^23^ to perform LC-MS/MS validation (level 1; according to metabolomics standard initiative). The spectrum represents the lactam form (2-hydroxy-5-oxo-proline) with three major fragmentation peaks (negative mode) at 126.0 Da, 82.0 Da and 42.0 Da (**Suppl. Fig. 1A**). αKGM was also detected in positive mode yielding a fragments of 105,1 and 91 Da (data not shown). Using pure αKGM as a standard, we confirmed that αKGM was increased in endothelial cells exposed to 300 µM H_2_O_2_ (**Suppl. Fig. 1B**). Furthermore, we determined the concentration of αKGM in plasma and urine from healthy individuals. The average plasma levels of αKGM and αKG were 3.4 and 12.3 µM, respectively. In urine, the concentrations were 19.6 and 9.3 µmol/mmol of creatinine for αKGM and αKG, respectively (**Suppl. Fig. 1C**).

### A single nucleotide variant (SNV) decreases NIT2 expression and elevates plasma levels of αKGM in humans

To determine a potential correlation between NIT2 and αKGM levels in humans, we performed a metabolite quantitative trait locus analysis (mQTL) along with an expression quantitative trait locus analysis (eQTL). For this analysis we used datasets^24^ generated by Metabolon^®^ and we confirmed the identity of αKGM using LC-MS/MS. One sentinel SNV (rs38380303, chr3:100334840:C/GC) showed a highly significant association with increased plasma αKGM (p-value 1.4×10^−36^)^24^. However, this SNV localizes in an indel and is not present in many genome-wide association studies. The second most significant SNV is rs277627 (chr3:100336429: G/A) that is in linkage distribution to rs38380303 and has a p-value of 2.0×10^−36^ for increased αKGM (**Fig. 2A**). rs277627 is located in intron 1 of the *NIT2* gene, which contains a regulatory element (REM, enhancer coordinates: chr3:100336401-100336500, hg38) that can potentially affect the binding of transcription factors (TF). Of the 15 transcription factors that bind to this regulatory element and are expressed in human endothelial cells (RNAseq data, **Suppl. Table 4**), we observed that the binding of 9 of them was likely to be affected by the mutation (**Suppl. Fig. 2A**). To verify this, we deleted the REM using CRISPR/cas9 in HEK 293 cells (**Fig. 2B-C, Suppl. Fig. 2B**). Deletion of the locus containing rs277627 resulted in a decrease in expression of NIT2 mRNA and protein (**Fig. 2D-E**). In line with this, the rs277627 GA and AA variants resulted in a lower expression of NIT2 in human aorta and tibial artery samples (GTEx data^25^, **Suppl. Fig. 2C**), than subjects carrying GG. Phenome-wide association studies to rs38380303 and rs277627 (**Suppl. table 1 and 2)** revealed a positive association of rs277627 with risk factors for hypertension (odds ratio: 1.06, p=0.01; CLSA cohort, analyzed with logistic regression. Fasting hour, sex, age, BMI, and recruitment centers adjusted in the model).^24^ Thus, a variant of the *NIT2* gene that leads to the accumulation of αKGM in the plasma, resulted in decreased NIT2 mRNA and protein levels in humans.

**Figure 2:**
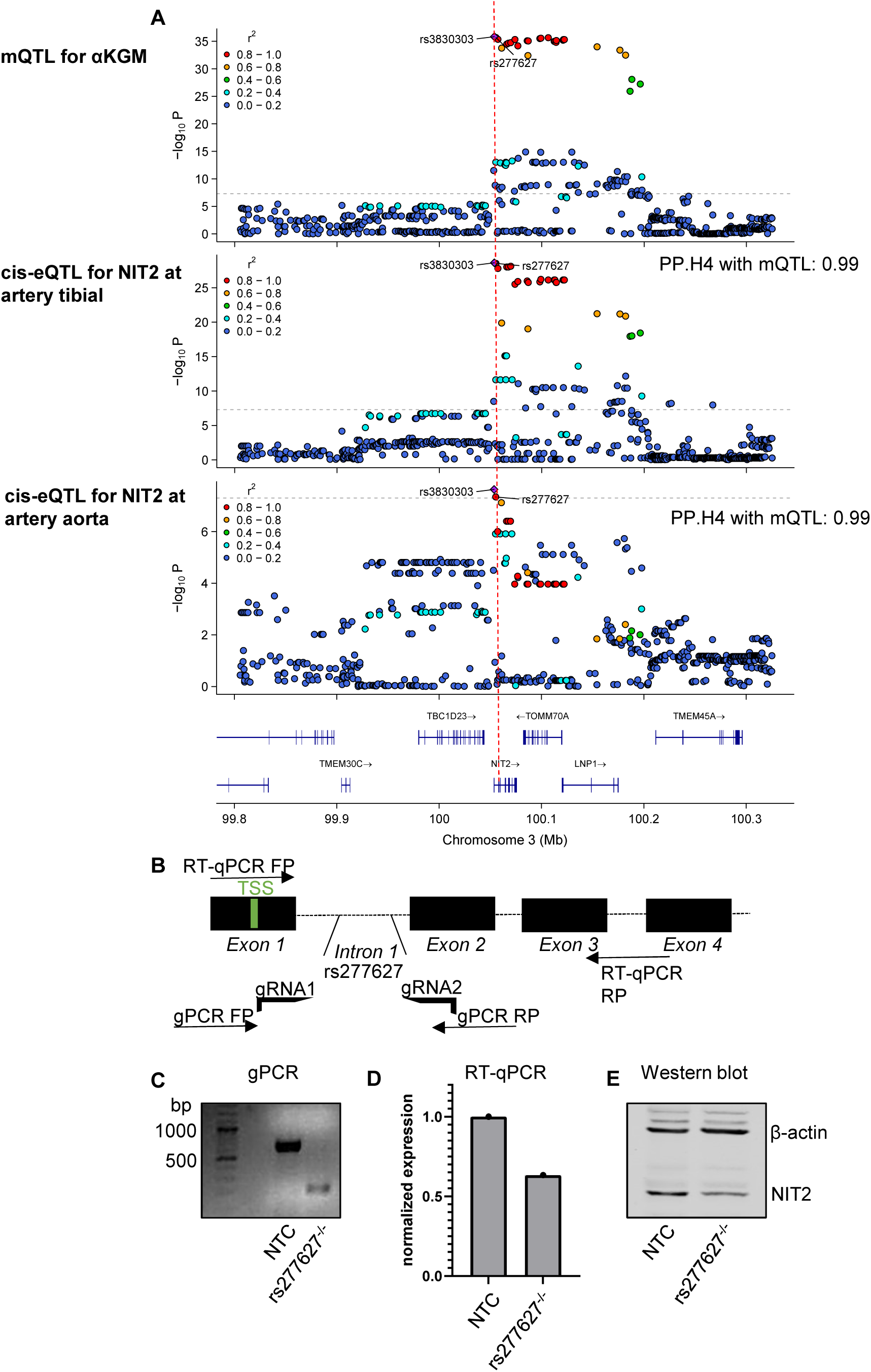
A human single nucleotide variant (SNV) decreases *NIT2* expression and elevates plasma levels of αKGM. A: Colocalization analyses and regional association plots of αKGM mQTL (CLSA metabolite, n = 8203) with cis-eQTL for *NIT2* in artery tibial (n = 475) and aortic tissue (n= 329) (GTEx v8 study). The sentinel variant rs3830303 and rs277627 are indicated. B: strategy for CRISPR/cas9 deletion of the regulatory element containing rs277627 in HEK 293 cells. C: Genomic PCR showing a 437 bp deletion of the rs277627 containing locus. *NIT2* expression in NTC and rs27767^−/−^ HEK 293 cells by RT-qPCR (D) and Western blot (E). TSS: transcription start site. RT-qPCR: Real-time polymerase chain reaction. FP: forward primer. RP: reverse primer. NTC: non-targeting control.

### H_2_O_2_ inactivates NIT2 by cysteine oxidation

Since lower NIT2 activity, either induced by exposure to H_2_O_2_ or due to decreased expression leads to the accumulation of αKGM, we determined whether αKGM is increased in response to a pro-inflammatory condition associated with oxidative stress *in vivo*. αKGM and αKG were determined in mice treated with vehicle or lipopolysaccharide (LPS, 4 mg/kg) for four hours. Plasma levels of neither metabolites were altered after LPS injection but urinary levels of αKGM were significantly increased and levels of αKG decreased (**Fig. 3A-B**). The observation that the differences were confined to urine can probably be attributed to a high clearance of αKGM in its lactam form or a high renal production.

**Figure 3:**
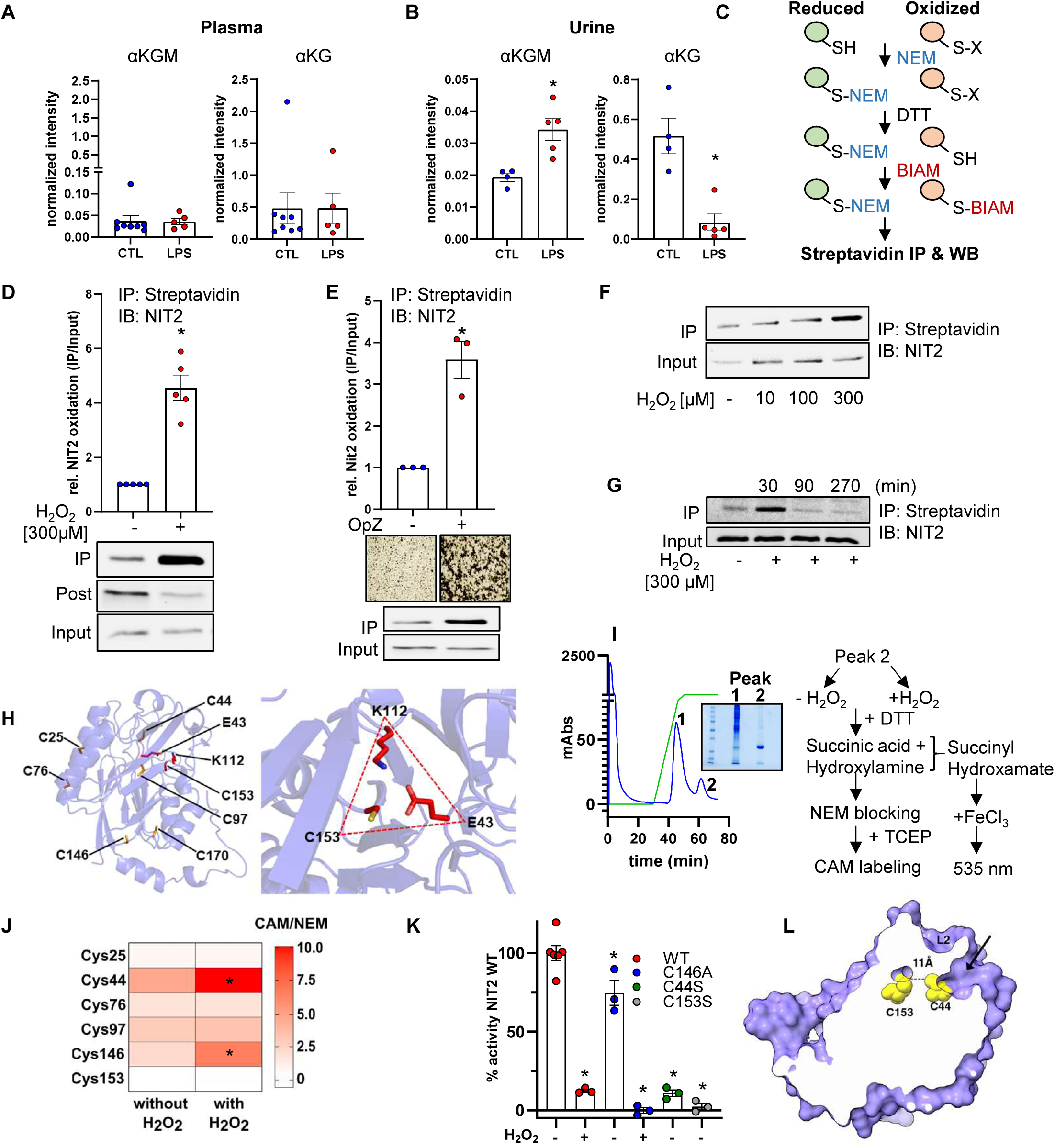
H_2_O_2_ inactivates NIT2 by cysteine oxidation. Targeted LC-MS/MS for αKGM and αKG in plasma (A) and urine (B) of mice treated with lipopolysaccharide (LPS, 4 mg/kg, 4 h). C: Schematic representation of biotinylated iodoacetamide (BIAM) switch assay. D-F: BIAM switch assay followed by immunoblotting for NIT2 in HUVEC (D) exposed to H_2_O_2_ or human granulocytes (E) incubated with or without zymosan opsonized by human plasma. *p<0.05, without *vs.* with stimulation, t-test. Microscopy images show aggregation of granulocytes in response to zymosan. F: BIAM switch assay with a H_2_O_2_ concentration and time (G) response curve. H: Cartoon representation of the predicted structure of human NIT2 (AlphaFold2). The 7 cysteine residues are depicted. The insert shows a zoom in view of Cys153 and the catalytic centre (C153-L112-E43). I: Schematic representation of overexpression, affinity purification, redox LC-MS and activity assay for NIT2. J: Heatmap summarizing reversible modifications of Cys in NIT2. K: NIT2 activity assay using competitive amine substitution of succinic acid by hydroxylamine. *p<0.05 as compared to wild type NIT2 without H_2_O_2_, one-way ANOVA with Bonferroni correction. L: Solvent exposed surface of the predicted NIT2 model depicted in purple. Cys153 and Cys44 (yellow spheres) are 11 Å apart. Cys44 localizes at the bottom of the open substrate binding channel capped by the flexible loop 2 (L2). Arrow shows a tunnel from the protein surface down to the Cys153. IP: immunoprecipitation. IB: immunoblotting.

To determine whether cysteine oxidation could contribute to the inhibition of NIT2 in pro-inflammatory conditions with increased ROS production, we used the biotinylated iodoacetamide (BIAM) switch assay^26^ (**Fig. 3C**). It was possible to demonstrate that H_2_O_2_ increased the cysteine oxidation of NIT2 (**Fig. 3D**). More importantly, a similar effect was observed when endothelial cells were co-incubated with human granulocytes, which were pre-activated with opsonized zymosan to stimulate ROS production (**Fig. 3E**). When cells were treated with H_2_O_2_, NIT2 oxidation was dependent on both H_2_O_2_ concentration and exposure time, with the largest effect occurring at 300 µM H_2_O_2_ and 15-30 minutes of exposure (**Fig. 3F-G, Suppl. 3A**). To determine whether NIT2 oxidation was restricted to endothelial cells, the BIAM switch assay was repeated using several other cell types including human carotid and aortic endothelial cells, fibroblasts, smooth muscle cells and HEK 293 cells. In all of the cell types studied, H_2_O_2_ elicited the oxidation of NIT2 (**Suppl. Fig. 3A-F**). However, the oxidation of NIT2 was not induced by other types of ROS as neither diamide (up to 100 µM) nor menadione (up to 50 µM) were able to oxidize NIT2 (**Suppl. Fig. 3G-H**).

NIT2 contains seven cysteine residues with one residing at the catalytic triad (C153-K112-E43, **Fig. 3H**). Interestingly, NIT2 in vertebrates has higher Cys content than that of other species and only Cys153 is conserved down to yeast, bacteria and plants (**Suppl. Fig. 4**). To identify the cysteine residues, which are oxidized by H_2_O_2_ we first performed LC-MS of the proteins enriched in the BIAM switch assay. NIT2 could be identified but with low peptide counts and insufficient coverage. Alternatively, NIT2 was pulled down using an anti-NIT2 antibody instead of the streptavidin antibody, but the NIT2 peptide count was too low for quantitative measurements and not all cysteines were identified. The only successful approach that identified multiple peptides covering six out of seven cysteine residues of NIT2 was based on the overexpression of a His-tagged NIT2 in HEK 293 followed by its purification by affinity chromatography. This approach yielded one peak (peak 2) containing pure NIT2 (**Fig. 3I**, see Coomassie blue staining). The purified enzyme was subsequently treated without or with H_2_O_2_ (300 µM) in the presence of succinic acid and hydroxylamine for the identification of the redox-sensitive cysteines and activity assay as depicted in **figure 3I**. Using this approach the ratio of chloroacetamide (CAM) to N-ethylmaleimide (NEM) labelling should reflect the reversible oxidations induced by H_2_O_2_. While the Cys153 was not significantly modified by H_2_O_2_, Cys44 and Cys146 showed a two and a three-fold significant increase in CAM/NEM ratio, respectively when exposed to H_2_O_2_ indicating a reversible oxidation of these cysteine residues (**Fig. 3J**).

To investigate how Cys modification affected NIT2 we assayed enzyme activity by following the competitive amine substitution of succinic acid by hydroxylamine, as previously described.^27^ Incubation of wild type NIT2 with 300 µM H_2_O_2_ inhibited its activity by 90 % whereas a catalytically dead mutant (C153S) was inactive (**Fig. 3K**). Since Cys44 and Cys146 were reversibly oxidized by H_2_O_2_ we generated mutants of these cysteines. Cys146 is located in a structural loop (**Fig. 3H**) and replacing it with either serine (C146S) or aspartate (C146D) resulted in a low expression and low yield in affinity purification. Replacing the cysteine with alanine (C146A) resulted in an unstable protein, as detected using a thermal shift assay (**Suppl. Fig. 5A-C**). The C146A NIT2 mutant retained 70 % of the basal activity of the wild type enzyme but remained sensitive to H_2_O_2._ In contrast, the mutation of Cys44 to serine (C44S) resulted in a pronounced loss of activity (**Fig. 3K**). Cys44 is located in a tunnel that extends from the surface of the protein to the catalytic center, forming the substrate channel (**Fig. 3L, Suppl. Fig. 6A-D**). While Cys44 is approximately 11 Å from Cys153, the surrounding negative charge makes it unlikely that these cysteines form a disulphide bond. Therefore, we conclude that Cys44, which forms the substrate channel, is the main redox-sensitive cysteine, and its oxidation inhibits NIT2 activity, probably by hindering substrate access to the active center.

### NIT2 and GLS1 synergistically maintain the endothelial metabolome

Having identified NIT2 as the redox-sensitive enzyme that generates αKG from glutamine, we set out to investigate its contribution for glutamine metabolism in conjunction with the endothelial default pathway involving GLS1. CRISPR/cas9 was used to generate endothelial cells lacking NIT2 (NIT2^−/−^) and GLS1 (GLS1^−/−^) alone or in combination (NIT2/GLS1^−/−^). Western blot analysis and immunofluorescence staining confirmed successful knockout (**Fig. 4A-B**). Furthermore, immunofluorescence suggested that GLS1 has a mitochondrial localization whereas NIT2 is distributed across the cell (**Fig. 4B)**.

**Figure 4:**
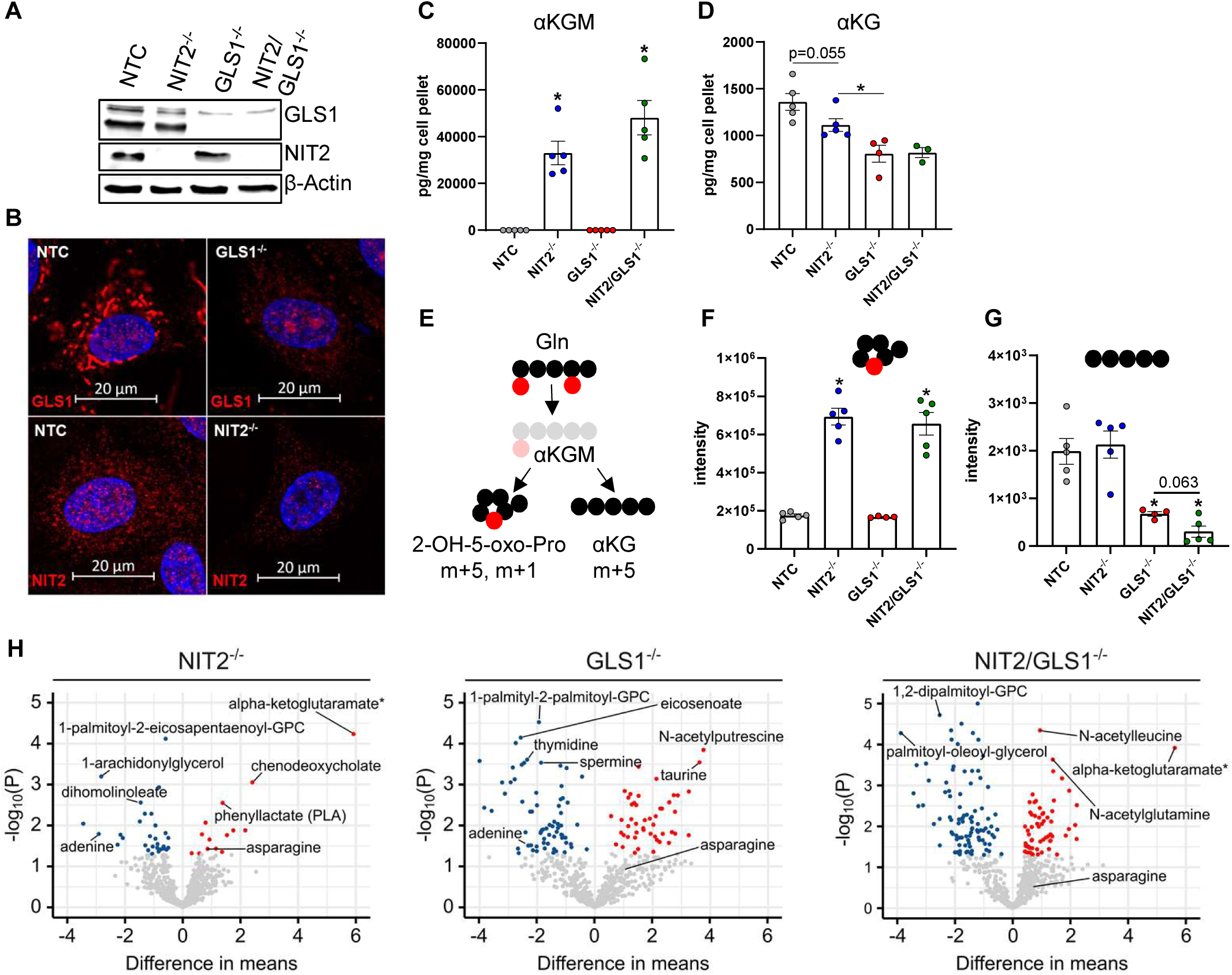
NIT2 and GLS1 synergistically maintain the endothelial metabolome. NIT2^−/−^, GLS1^−/−^ and NIT2/GLS1^−/−^ HUVEC were generated by CRISPR/cas9 and knockout efficiency was validated by Western blot (A). Cellular distribution of NIT2 and GLS1 as shown by immunofluorescence (B). C-D: Targeted LC-MS/MS measurements for αKGM and αKG in CRISPR/cas9 HUVEC. *p<0.05 as compared to NTC, ANOVA with Bonferroni correction. E: Scheme for isotopic tracing of fully labelled glutamine that generates m+5, m+1 αKGM and m+5 αKG. F: m+5, m+1 αKGM and m+5 αKG (G) in HUVEC. *p<0.05 as compared to NTC. H: Volcano plots of significantly altered metabolites as measured by untargeted metabolomics. NTC: non-targeting control.

To monitor the levels of metabolites, targeted LC-MS/MS measurements were performed. αKGM was not detected in either endothelial cells treated with a non-targeted construct (NTC) or in cells lacking GLS1, presumably as a result of high NIT2 activity. However, the deletion of NIT2 resulted in a marked increase in αKGM levels (**Fig. 4C**), accompanied by a slight decrease in αKG, which was more markedly affected by the deletion of GLS1 (**Fig. 4D**). To analyze glutamine utilization by the two pathways, we performed isotopic tracing with fully labeled glutamine (^13^C_5_, ^15^N_2_) and utilized a heavy isotope of αKGM (m+5, m+1) as reference (**Fig. 4E**). The knockout of NIT2 increased the production of m+5, m+1 αKGM by 4 fold (**Fig. 4F**) but did not affect the production of m+5 αKG. In contrast, deletion of GLS1 had no impact on m+5, m+1 αKGM levels but largely contributed to m+5 αKG (**Fig. 4G**). Importantly, the double knockout of NIT2 and GLS1 further decreased m+5 αKG as compared to the single GLS1 knockout. These observations suggest that despite of the large contribution of GLS1 to αKG levels in endothelial cells, NIT2 functionally generates αKG from glutamine. NIT2-derived αKG production seems to be particularly relevant when GLS1 activity is low.

To explore the overall contribution of NIT2 and GLS1 to the endothelial cell metabolome, we performed untargeted metabolomics of single and double knockout cells. αKGM stood out as the most significantly increased metabolite of those measured in NIT2^−/−^ and NIT2/GLS1^−/−^ cells (**Fig. 4H**). The deletion of NIT2 significantly altered 10 unique metabolites whereas knockout of GLS1 altered 51 metabolites. However, the combined deletion of both enzymes significantly altered 86 unique metabolites, suggesting that both pathways synergistically contribute to the metabolome of endothelial cells (**Suppl. Table 3**). A similar trend was observed regarding gene expression as the deletion of NIT2 differentially regulated 7 unique genes whereas GLS1^−/−^ altered 91 and NIT2/GLS1^−/−^ 1161 (**Suppl. Fig. 7, Suppl. table 4**).

Altogether, deletion of NIT2 led to an accumulation of αKGM and a decrease in αKG levels. GLS1 has a greater contribution to αKG production than NIT2 but both pathways are active and maintain the metabolome and gene signature of endothelial cells.

### Endothelial knockout of NIT2 impairs angiogenesis in mice

To explore the function of NIT2 *in vivo* we generated endothelial cell specific, tamoxifen-inducible knockout mice of NIT2 (CTL and ecNit2^−/−^, **Suppl. Fig. 8A-B**) as well as global, constitutive knockout mice (Nit2^wt/wt^ and Nit2^ko/ko^, **Suppl. Fig. 8C**). Efficient knockout of NIT2 in ecNit2^−/−^ was confirmed by Western blot of endothelial cells enriched from the aorta (**Fig. 5A**) and immunofluorescence in the retina vasculature (**Fig. 5B**). To evaluate the contribution of endothelial NIT2 to the glutamine system in mice, untargeted metabolomics was performed from plasma and lung tissue, which is rich in endothelial cells. In plasma, there was a trend towards increased αKGM (p=0.0506) in ecNit2^−/−^ as compared to control mice. Importantly, αKG was significantly decreased in ecNit2^−/−^ as compared to CTL mice, demonstrating a contribution of endothelial NIT2 to plasma αKG level (**Fig. 5C-E, Suppl. Table 5**). Untargeted metabolomics from whole lung showed αKGM as the second most upregulated metabolite in ecNit2^−/−^ as compared to CTL mice and although αKG was unchanged, there was a significant increase in glutamine level suggesting either substrate accumulation in the pathway or compensation (**Fig. 5F, Suppl. Table 6**). Altogether, metabolomics of ecNIT2^−/−^ mice suggest a contribution of the endothelium to cellular and systemic αKGM and αKG. Moreover, the knockout of NIT2 in mice affects metabolites outside of the GTωA pathway.

**Figure 5:**
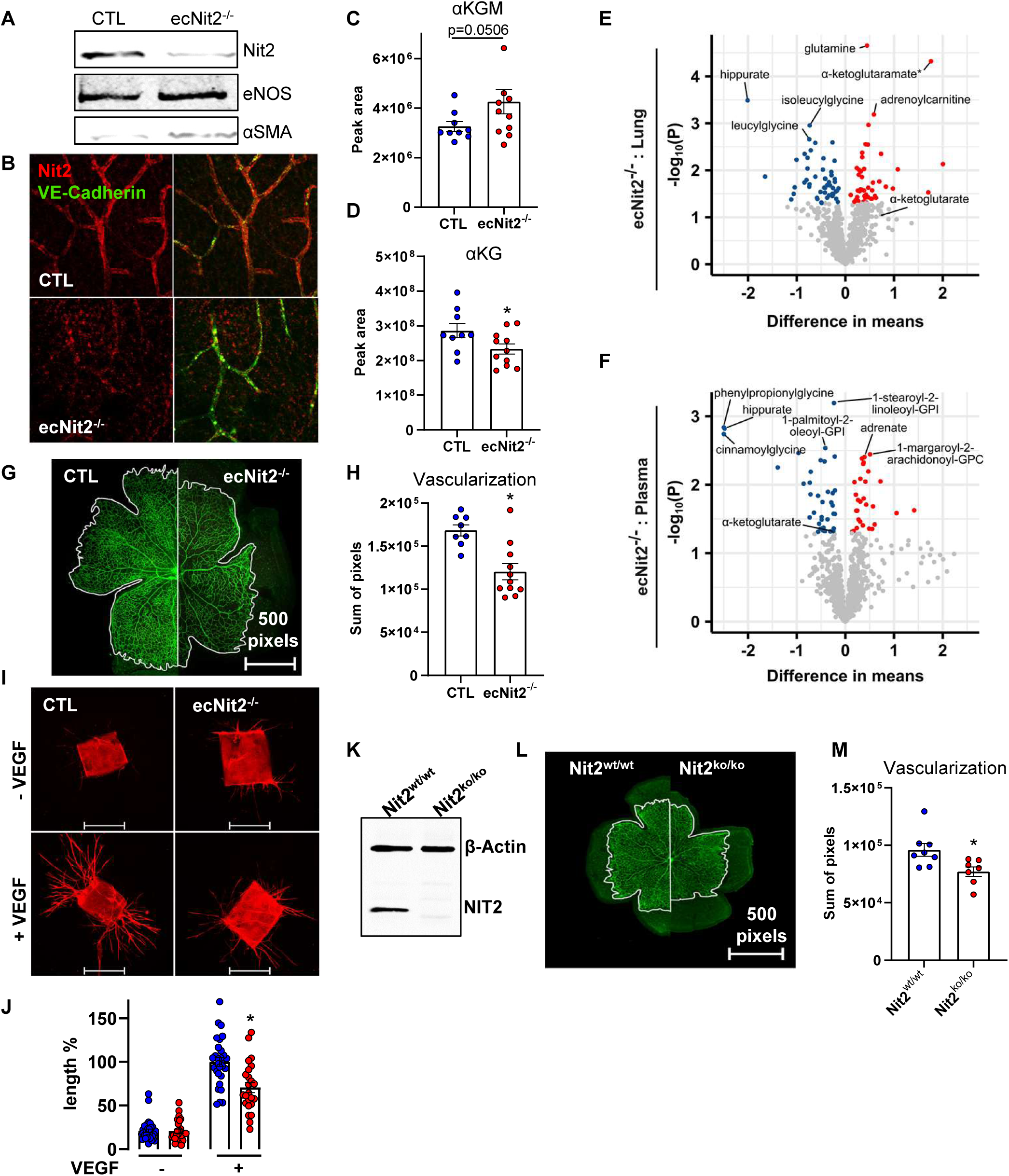
Endothelial knockout of NIT2 impairs angiogenesis in mice. Generation of a tamoxifen-inducible, endothelial cell-specific knockout mouse of Nit2 (ecNit2^−/−^). A: Validation of the knockout efficiency by Western blot of aortic endothelial cells and immunofluorescence in retina vessels (B). C-D: LC-MS/MS for αKGM and αKG in plasma of CTL and ecNit2^−/−^ mice. *p<0.05 as compared to CTL, Mann Whitney test. E-F: Untargeted metabolomics from CTL and ecNit2^−/−^ mice from lung tissue (E) and plasma (F). G: Retina angiogenesis in neonatal mice as indicated (post natal day 7), isolectin staining from CTL and ecNit2^−/−^ mice with quantification of vascularization (H). I: *Ex vivo* endothelial cell outsprouting from aortic segments isolated from CTL and ecNit2^−/−^ mice with quantification, normalized to CTL –VEGF (J). *p<0.05, Mann Whitney test (-VEGF versus +VEGF). K: Validation of NIT2 deletion by Western blot in aortic endothelial cells of a global, constitutive knockout mouse of Nit2 (Nit2^wt/wt^ and Nit2^ko/ko^). L: Retina angiogenesis in neonatal mice as in G with quantification (M). *p<0.05 Nit2^wt/wt^ as compared to Nit2^ko/ko^, Mann Whitney test.

For endothelial GLS1 an important contribution to angiogenesis is established.^15^ Thus, we examined whether deletion of NIT2 leads to a similar phenotype using the neonatal retina model of angiogenesis. Endothelial-specific knockout of NIT2 led to a decrease of vessel length by 30% as compared to retinae of CTL mice (**Fig. 5G-H**). Furthermore, in the aortic outgrowth assay (*ex vivo* model of angiogenesis), VEGF-induced sprouting was significantly decreased in ecNit2^−/−^ as compared to CTL mice (**Fig. 5I-J**). Importantly, also global knockout mice of Nit2 (Nit2^ko/ko^, which do not express Cre recombinase) exhibited decreased retinal angiogenesis, excluding any toxic effects of Cre recombinase activity^28^ (**Fig. 5K-M)**. In conclusion, endothelial knockout of NIT2 decreases angiogenesis similarly to endothelial deletion of GLS1.

### Endothelial deletion of NIT2 depletes adenine and induces senescence

To identify the mechanism underlying the attenuated angiogenic function of ecNIT2^−/−^ mice, functional experiments were performed with HUVEC. First, we recapitulated the angiogenic role of NIT2 using the spheroid outgrowth assay. NIT2^−/−^, similarly to GLS1^−/−^ and NIT2/GLS1^−/^, impaired endothelial cell sprouting under basal but also the VEGF-A-induced sprouting (**Fig. 6A-C**). Given that angiogenic function in this assay is a consequence of proliferative as well as migratory capacity of the endothelial cells, these aspects were differentiated. Neither knockout of NIT2 nor pharmacologic inhibition of GLS1 affected endothelial cell migration in the scratch wound assay (**Fig. 6D**). In contrast to this, both interventions decreased proliferation (**Fig. 6E)** with the combination of NIT2 knockout and GLS1 inhibition having an additive effect. Impaired proliferation with maintained migration may imply that synthetic function of the cells is attenuated. This could be a consequence of a lack of anaplerotic TCA cycle equivalents. However, it has been previously reported that decreased proliferation in glutamine-deprived endothelial cells was only partially rescued by adding extra TCA carbons in the form of cell-permeable compounds (dimethyl-αKG, monomethyl-succinate, oxaloacetate or pyruvate).^15^

**Figure 6:**
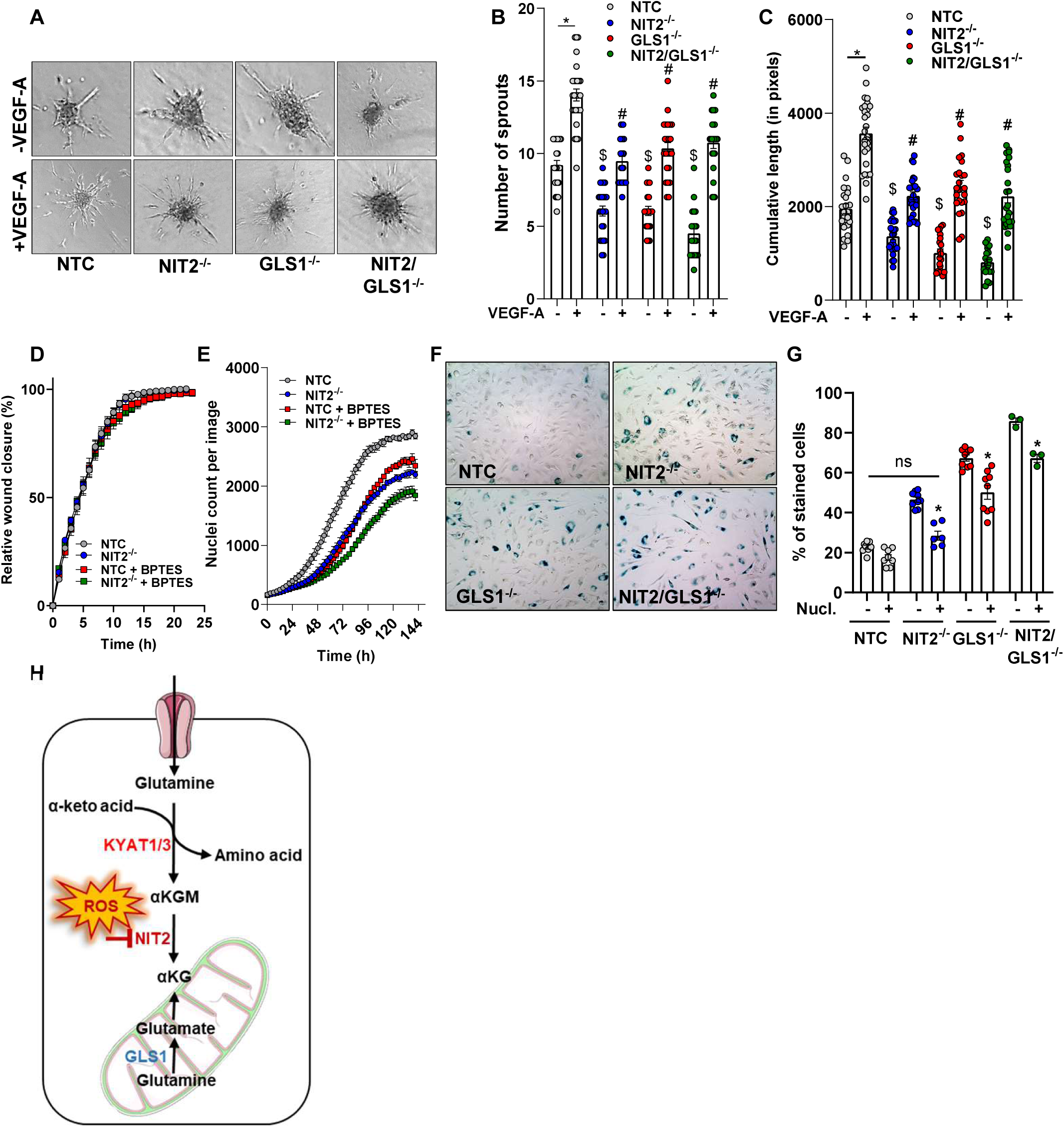
Deletion of NIT2 results in endothelial cell senescence. A-C: sprout outgrowth assay of HUVEC with and without VEGF-A (10 ng/mL) as indicated. $ p<0.05 as compared to NTC without VEGF-A and #p<0.05 as compared to NTC with VEGF-A, one-way ANOVA, Bonferroni correction. D: migration and (E) proliferation assays. F-G: senescence assay with β-galactosidase staining in HUVEC as indicated. Nucl: nucleosides. * p<0.05 as compared to NTC, one-way ANOVA with Bonferroni correction. H: NIT2 is a redox switch in the glutamine transaminase-ω-amidase pathway.

Interestingly, adenine level was significantly lower in endothelial cells knockout of NIT2 or GLS1, pointing to alterations in purine handling. Lack of nucleotides decrease proliferation and pushes cells towards senescence.^29^ To address whether a similar effect is operative in the present model, β-galactosidase staining was performed. Deletion of NIT2 or GLS1 both increased the number of senescent cells and combined deletion of both enzymes resulted in an additive effect (**Fig. 6F**). In order to determine whether this effect was a consequence of a potential shortage of nucleotides, a nucleoside mix (EmbryoMax^®^ Nucleosides, Merck #ES-008) was administered. This reverted the NIT2-deletion induced senescence phenotype and only attenuated senescence rates in GLS1^−/−^ and double knockout cells (**Fig. 6G**).

## Discussion

In this study, we aimed to identify metabolic targets of oxidative stress in endothelial cells. For this, we exposed HUVEC to menadione or H_2_O_2_ as oxidizing agents that have different properties. Exposure of endothelial cells to H_2_O_2_ selectively unmasked a metabolic pathway for glutamine that has been largely overlooked and underappreciated. GLS1 within the glutaminase I pathway was hitherto the only known enzyme of glutamine metabolism contributing to endothelial αKG production. Here we present the glutamine transaminase-ω-amidase (GTωA) pathway that is a redox switch in glutamine metabolism and important for non-canonical production of αKG and angiogenesis.

The GTωA pathway consists of two coupled reactions: i. the transamination of glutamine by the enzymes KYAT1 and KYAT3 to generate αKGM. This reaction is accompanied by the replenishment of amino acids using their α-keto acids as substrate. ii. αKGM is converted to αKG by NIT2. Exposure of endothelial cells to H_2_O_2_ increased αKGM and conversely decreased αKG, suggesting that the reaction catalyzed by NIT2 is redox sensitive. Using BIAM switch assay, redox proteomics and site directed mutagenesis we demonstrated that NIT2 is reversibly oxidized by H_2_O_2_ and that Cys44 and Cys146 are the reactive cysteines. Cys146 is located in a surface loop and scores the highest with algorithms that predict cysteine oxidation.^30^ However, there is no evidence nor modeling that predict functional consequences like conformational changes upon oxidation of Cys146. This cysteine seems to be rather important for proper folding or stability of the protein, as the C146S and C146D mutants were unstable. In contrast, Cys44 was directly linked to the catalytic activity of NIT2. Despite of the high confidence score for the modeling (> 90) with AlphaFold2 for human NIT2, its crystal structure is not available and the mouse structure^31^ does not contain the substrate bound to it what limits the understanding of the role of Cys44 in NIT2 catalysis. The variant C44S abolished NIT2 activity, suggesting that given its localization in the open substrate tunnel, Cys44 is the primary redox switch and it might have an important role in the initiation of of the catalytic cycle.

How relevant is NIT2 for endothelial metabolism? To address this question, we focused on αKG, which is a major metabolic hub. Knockdown of GLS1 decreases αKG by 40%^14^, so that alternative pathways are relevant. This could be via import of αKG from the medium through transporters^32^, the formation of αKG in the TCA cycle and the NIT2 ω-amidase reaction. Measurements of cellular αKG content and isotopic tracing of glutamine suggested that the GLS1 pathway is dominant for αKG. NIT2, however, contributes to about 20% of the αKG formation from glutamine as well as to the total αKG pool. The *in vivo* importance of endothelial NIT2 for αKGM and αKG pools was studied by generating ecNit2^−/−^ mice. Endothelial-specific deletion of NIT2 led to an accumulation of αKGM in plasma whereas αKG levels were conversely decreased. Given that all cells, but particularly liver and kidney are thought to contribute to plasma αKG, the latter finding is unexpected. It demonstrates that *in vivo* under normal conditions, the relevance of endothelial NIT2 for αKG formation might be greater than what is suggested by the cell culture studies.

Functionally, the knockout of NIT2 in endothelial cells had similar effects to that of the knockout of GLS1^14,15^: it decreased αKG pools, did not affect migration but decreased proliferation and decreased angiogenesis in endothelial cells and in the mouse retina. Moreover, deletion of NIT2 resulted in premature senescence of endothelial cells in culture. Double knockout of NIT2 and GLS1 (NIT2/GLS1^−/−^) potentiated all these effects, which suggest that both glutamine pathways compensate each other. The question that therefore arises is why endothelial cells do contain both pathways? The GTωA may be advantageous for endothelial cells as the production of αKG from glutamine does not involve a net oxidation and thus does not regenerate NAD(P)H as in the glutaminase I pathway. In this way, αKG production can be uncoupled from mitochondria. Indeed, we observed by immunofluorescence that NIT2 is distributed across endothelial cells and not restricted to the mitochondria as GLS1. The fact that NIT2 is redox sensitive might be a way to block the entrance of αKG into the mitochondria what could potentiate ROS production and oxidative stress. Another potential metabolic advantage of GTωA pathway might be the salvage of α-keto acids to their corresponding amino acids (**Fig. 6H**).^33^

It is surprising that the GTωA pathway has so far not gained much attention. Potentially, this is a consequence of lack of standard, but also of the short half-life of αKGM in its linear form and its effective de-amidation to αKG by NIT2. αKGM (lactam form) levels in cells are essentially undetectable in the presence of active NIT2 and plasma levels are lower than αKG, as demonstrated in the present study. αKGM was previously produced through oxidation of L-glutamine with snake venom L-amino acid oxidase in the presence of catalase.^23^ αKGM measurements in its 2-hydroxy-5-oxo-proline form were originally performed by gas chromatography or HPLC.^18^ Such methods have drawbacks but set the basis for the first characterization of the GTωA pathway under normal and pathological conditions. For example, αKGM is elevated in the cerebrospinal fluid of hyperammonemic patients with hepatic encephalopathy and in the urine of hyperammonemic patients with an inborn error of urea cycle or citrin deficiency.^34–38^

The detection and semi-quantification of αKGM in untargeted metabolomics was based on the predicted fragmentation pattern. The identity of αKGM was validated for the first time in this study via LC-MS/MS. Importantly, data from untargeted metabolomics that contain αKGM^24^ allowed for a co-localization analysis that found an association of a NIT2 variant with higher αKGM in plasma. Additionally, the increasing number of publicly available metabolomic datasets provides opportunities to mine data for associations between αKGM and diseases, and to further evaluate the functions of the GTωA pathway. This applies in particular to cells and tissues with high glutamine utilization, such as cancer cells, the liver, and the kidneys. From metabolomics datasets available, αKGM is one out of nine significantly increased metabolites in serum of patients with chronic and large ischemic infarction volume.^39^ In patients with chronic kidney disease αKGM is among the five significantly increased metabolites with the strongest associations with urinary uromodulin.^40^ Inasmuch as αKGM is increased in urine of LPS-treated mice, it is possible that this metabolite is a marker for oxidative stress in pro-inflammatory conditions as a result of NIT2 inhibition under oxidative conditions.

In conclusion, we present the ω-amidase/NIT2 as a molecular target of oxidative stress and a redox-switch in metabolism. Its inhibition by oxidation at cysteine residues leads to an accumulation of αKGM, a metabolite that was previously not fully studied. Our work provides the first systematic analysis of the role of NIT2 in the GTωA pathway, which serves as an alternative and non-canonical route for the generation of αKG from glutamine in endothelial cells. Deletion of NIT2 decreased endothelial αKG pool, cell proliferation, angiogenesis and induced senescence. Our findings in EC can certainly inspire further research into the general glutamine metabolism and the further biological importance of αKGM and the GTωA pathway.

### Limitations of this study

1. The lack of a crystal structure for human NIT2 with the substrate bound at its active site limited our ability to model how oxidation of Cys44 and Cys146 affect conformational changes, substrate binding, and multimerization that may be important for NIT2 activity.
2. We focused on the endothelial function of NIT2 due to the dependence of endothelial cells on glutamine and our scientific interest and expertise. However, the GTωA pathway is predicted to be important also in other cell types and organs, particularly those with high glutamine utilization, such as many cancer cells, the liver, and the kidneys.
3. We observed lower levels of adenine in NIT2^−/−^ endothelial cells as compared to control. The direct link between NIT2 and adenine, however, is unclear from the so far established metabolic pathways.

## Supporting information

Phenome-wide association studies to rs38380303

Phenome-wide association studies to rs277627

Untargeted metabolomics of HUVEC NTC, NIT2-/-, GLS1-/- and NIT2/GLS1-/-

Differential gene expression analysis (RNAseq) of HUVEC NTC, NIT2-/-, GLS1-/- and NIT2/GLS1-/-

Untargeted metabolomics of plasma from CTL and ecNit2-/- mice

Untargeted metabolomics of lung tissue from CTL and ecNit2-/- mice

## Acknowledgments

We dedicate this study to Prof. Arthur J.L. Cooper (* 04.19.1946 † 05.30.2024) who was emeritus professor at Medical College in Valhalla, NY and adjunct professor at Weill Cornell Medical College, NY, United States. He studied NIT2 and the GTωA pathway in the 1970s and had a substantial intellectual contribution to the present study.

We are grateful for excellent technical assistance of Susanne Wienströer, Katalin Pálfi, Jana Meisterknecht, Manuela Späth and Natalie Weber.

## Source of funding

The study was supported by the Deutsche Forschungsgemeinschaft (DFG): SFB 1531 – Project number 456687919 to R.P.B., F.R., I.F., M.S. and I.W. Germany‘s Excellence Strategy CIBSS – EXC-2189 – Project ID 390939984 to LH. Excellent Cluster Cardio-Pulmonary Institute (EXS2026, project number 390649896), the Medicine Faculty of the Goethe University (Frankfurt, Germany); the German Center for Cardiovascular Research (DZHK), Partner Site Rhein-Main, Frankfurt am Main.

## Disclosures

The authors declare no conflict of interest that could potentially influence or bias the work. T.D. co-owns the LiT Biosciences.

J.B.R. is the CEO of 5 Prime Sciences (www.5primesciences.com), which provides research services for biotech, pharma, and venture capital companies for projects unrelated to this research. He has served as an advisor to GlaxoSmithKline and Deerfield Capital. J.B.R.’s institution has received investigator-initiated grant funding from Eli Lilly, GlaxoSmithKline, and Biogen for projects unrelated to this research. Y.C. is an employee of 5 Prime Sciences.

J.T. has received speaking and/or consulting fees from Astra-Zeneca, Gore, Boehringer-Ingelheim, Falk, Grifols, Genfit and CSL Behring.

## Supplemental Materials

Excel table 1: Phenome-wide association studies to rs38380303

Excel table 2: Phenome-wide association studies to rs277627

Excel table 3: Untargeted metabolomics of HUVEC NTC, NIT2^−/−^, GLS1^−/−^ and NIT2/GLS1^−/−^

Excel table 4: Differential gene expression analysis (RNAseq) of HUVEC NTC, NIT2^−/−^, GLS1^−/−^ and NIT2/GLS1^−/−^

Excel table 5: Untargeted metabolomics of plasma from CTL and ecNit2-/-mice

Excel table 6: Untargeted metabolomics of lung tissue from CTL and ecNit2^−/−^ mice

## Methods

Protocols will be fulfilled by Dr. Flávia Rezende. Mouse lines will be shared under a Materials Transfer Agreement. Results from untargeted metabolomics and RNAseq are available in the supplementary material.

### Cell culture

Pooled human umbilical vein endothelial cells (HUVEC) were purchased from PromoCell (#C12203, Heidelberg, Germany) and cultured in endothelial growth medium (EGM), consisting of endothelial basal medium (EBM) supplemented with 8 % fetal calf serum, 0.5 % penicillin/ streptomycin (50 µg/mL), growth factors (EGF, bFGF, IGF, VEGF, #PB-C-MH-100-2199, PeloBiotech, Germany), heparin, L-glutamine but without hydrocortisone. For each experiment, at least three different batches of HUVEC at highest passage number 4 were used. Human embryonic kidney 293 cells (HEK 293) (ATCC, Manassas, USA) and Lenti-X 293 T cells (Takara, 632180, Japan) were cultured in Dulbecco’s Modified Eagle Medium High Glucose (Gibco) supplemented with 8% FCS, penicillin/ streptomycin (50 μg/mL of each) (#15140-122, Gibco/ Lifetechnologies, USA). All cells were cultured in a humidified atmosphere (5% CO_2_, 37 °C).

### Untargeted metabolomics and data analysis

Untargeted metabolomics of HUVEC exposed to menadione (5 μM) or H_2_O_2_ (300 μM) has been previously published.^16^ Untargeted metabolomics of CRISPR/cas9 NIT2^−/−^, GLS1^−/−^, and NIT2/GLS1^−/−^ HUVEC was performed from cells grown in EGM as described.^16^

Batch-normalized intensity data were analyzed for differential metabolite levels in R (https://ropensci.org/) using the metabodiff (v0.9.5) package. The data were subjected to k-nearest neighbor imputation, with a cutoff value of 0.25. Further normalization was performed using variance-stabilizing normalization, followed by quality checks to ensure no artefacts were introduced by the procedure. Corresponding treatment and control samples were then compared using the differential test function, which implements a Student’s T-Test followed by Benjamini-Hochberg correction for multiple testing. Results were visualized using ggplot2 (v3.3.5, RRID:SCR_014601).

### Establishment of αKGM as LC-MS/MS standard

Pure αKGM was synthesized as previously described.^23^ The heavy isotopologue (m+5 for carbons and m+1 for nitrogen) of αKGM was synthesized from [^13^C_5_-^15^N_2_]-L-Glutamine (Cambridge Isotopes Laboratories Inc., Tewksbury, USA). The fragmentation of αKGM was evaluated on a QTrap 5500 LC-MS/MS mass spectrometer (Sciex) via direct injection. Different collision energies (CE) were first tested in negative ionization mode, and a CE of 40 gave the best fragmentation pattern of m/z 144 Da (parental ion) as m/z 126 Da (most abundant), 82 Da (2^nd^ most abundant) and 42 Da (3^rd^ most abundant). Alternatively, αKGM was measured in positive ionization mode using a Sciex 6500+ ESI-tripleQ MS/MS on low mass mode (0–1000 Da) with declustering potential, exit potential, collision energy and collision cell exit potential of 206, 10, 21, 12 and 206, 10, 25 and 14 Volts, respectively. The dwell time was 20 milliseconds. Two fragments of the parental ion of αKGM (m/z 147 Da) showed m/z 105,1 Da and m/z 91 Da.

### Metabololite Quantitative Trait Locus (mQTL) in NIT2

To evaluate whether the same genetic variants are driving the associations with circulating αKGM levels and NIT2 transcript levels, a co-localization analysis was employed. The genetic associations for plasma αKGM nearby NIT2 coding region were obtained from Chen et al. (2023).^24^ Their genome-wide association study (GWAS) was conducted using the genetic data and metabolite measurements of individuals in European ancestry from the CLSA (Canadian Longitudinal Study on Aging) cohort (n = 8203). The *cis*-eQTL summary statistics for NIT2 in artery tibial and artery aorta were obtained from the GTEx v8 (n = 584 for artery tibial and n= 387 for artery aorta).^25^ Specifically, we applied a stringent Bayesian method using coloc R package (5.1.0) with the priors recommended by the original study *(P*1 = 1 × 10^−4^, *P*2 = 1 × 10^−4^, *P*12 = 1 × 10^−5^)^41^ to estimate the posterior probabilities (PP) that the αKGM and the NIT2 transcript share a single causal genetic locus. The genetic variants in the ±500 kb window of the leading metabolite quantitative trait locus (mQTL) of αKGM, rs3830303, that has minor allele frequency (MAF) over 0.05 were used for the analysis. The PP that two traits share one causal SNV (single nucleotide variant) above 80 %, namely PP.H4 > 0.8, was considered to be colocalized. LocusZoom plots for the tested genetic region were plotted using locuszoomr R package (0.1.3). The linkage distribution (LD) r^2^ for the genetic variants was obtained from the genetic data of European individuals in the 1000 Genome phase 3 reference panel.^42^ Genetic variants were plotted based on their LD to the leading mQTL. The αKGM GWAS from individuals of European ancestry can be found on the GWAS Catalog (https://www.ebi.ac.uk/gwas/) with accession number (GCST90199885). GTEx V8 release of eQTL data from individuals of European ancestry can be found in https://www.gtexportal.org/home/datasets.

### rs277627 analysis

To identify whether the SNV rs277627 is regulatory, we applied SNEEP (v1.0, 10.5281/zenodo.10830008), a statistical approach to identify whether a SNV significantly affects a TF binding site.^43^ For rs277627 (chr3:100336428-100336429, hg38) we compared the risk allele A against the major allele G. SNEEP requires as input a TF motif set, for which we used 817 non-redundant human motifs from JASPAR (version 2022) (10.1093/nar/gkab1113), HOCOMOCO (10.1093/nar/gkx1106) and the Kellis ENCODE motif database (10.1093/nar/gkt1249). To link the SNV to potential target genes, we downloaded 2.4 million regulatory elements (REM) associated to target genes, which are publicly available at the EpiRegio webserver (https://doi.org/10.1093/nar/gkaa382,REM-gene interactions: 10.5281/zenodo.3750929, filename: REMAnnotation.txt). The following parameters were specified to run SNEEP: -p 0.5, -c 0.01, -r REMAnnotation.txt, -g ensemblID_GeneName.txt. The SNEEP software, considered TF motifs, the file ensemblID_GeneName.txt and more details about the specified parameters are available at the GitHub repository (https://github.com/SchulzLab/SNEEP).

### Deletion of the regulatory element (REM) in the NIT2 gene

The REM that contains the SNV rs277627 was deleted by CRISPR/cas9 in HEK 293 cells using a dual gRNA approach (see CRISPR/cas9 section). After viral transduction, cells were clonally expanded by serial dilution and single clones were obtained. Validation of the CRISPR/Cas9 knockout of the target REM was performed by genomic PCR and Sanger sequencing from genomic DNA. CRISPR/Cas9 target sites were amplified by PCR (forward primer: ATCAGAGGCCAGGGTTTGTC, reverse primer: GAGGAAGGCAGGCATCTGTC) with PCR Mastermix (ThermoFisher, #K0171) and 500 ng DNA followed by agarose gel electrophoresis and ethidium bromide staining. Bands were excised from the gel and subjected to Sanger sequencing to further validate a deletion of 437 bp.

### RT-qPCR

Total RNA isolation was performed with the RNA Mini Kit (Bio&Sell) according to the manufacturers protocol and reverse transcription was performed with the SuperScript III Reverse Transcriptase (#12574026, Thermo Fisher Scientific, Massachusetts, USA) using a combination of oligo(dT)23 and random hexamer primers (Sigma). The resulting cDNA was amplifed in an AriaMX cycler (Agilent) with the ITaq Universal SYBR Green Supermix and ROX as reference dye (Bio-Rad, #1725125). Forward primer: GGTGCTTGTCTGCAGAGTCAT, reverse primer: TCTTCAGGGATAGAGCCTCCAA for *NIT2*. Relative expression was calculated using the ΔΔCt method and genes were normalized to β-Actin.

### Biotinylated iodoacetamide (BIAM) switch assay

HUVEC were exposed to either H_2_O_2_ (300 µM), menadione (5 or 50 µM) or diamide (100 or 500 µM) in EBM in a time course manner. Thiols were blocked with N-ethylmaleimide (NEM, 100 mM) and BIAM switch assay was performed as previously described.^26^

### Expression and purification recombinant of NIT2

The human 10x His-Tag NIT2 plasmid was obtained from Sino Biological (#HG23517-CH, Wayne, USA). NIT2 mutants were generated by site-directed mutagenesis to obtain NIT2-C146S, C146D, C146A and C44S (Q5^®^ Site-Directed Mutagenesis Kit, New England Bio Labs # E0554S).

Plasmid was transiently overexpressed for 48 hours in HEK 293 cells using PEI (polyethyleneimine, DNA: PEI ratio 1:5) for transfection. Subsequently, cells were lysed in Triton X-100 buffer supplemented with protease inhibitors. The extract was loaded on a HisTrap FF column packed with Ni-sepharose (Cytiva, #17531901) and purified using an Äkta FPLC system (GE Healthcare/ Cytiva, Solingen, Germany) with a flow rate of 0.5 mL/min. Proteins were eluted with a linear gradient of imidazole up to 500 mM.

### Thermal shift assay

Thermal shift assay was performed as previously described.^44,45^ Briefly, differential scanning fluorimetry was performed in a PCR plate with a total volume of 40 µL. Purified NIT2-WT (*C*_final_ 5 µM) or NIT2-C146A (*C*_final_ 5 µM), Triton X-100 (0.001% w/v), and SYPRO Orange (Thermo Fisher Scientific) (2.5x) were mixed in phosphate buffer with or without DTT (100 mM KPO_4_, 5 mM DTT). The samples were measured on an Icycler IQ single-color real time PCR system (λ_ex_ = 490 nm, λ_em_ = 570 nm) and emission was recorded during a temperature gradient of 0.2 °C increase per 24 s (25 – 80 °C). Raw data from both, NIT2-WT and NIT2-C146A, measurements were analyzed directly using a Boltzmann sigmoidal fit in GraphPad Prism. The *V_50_* values were considered as the melting temperatures. All samples were measured in triplicates.

### NIT2 activity assay

To measure NIT2 activity, the wild type and mutants of NIT2 were treated with H_2_O_2_ (300 µM) for ten minutes in 100 mM KPO_4_ buffer, pH 7.4. Next, samples were incubated with 5 mM DTT for ten minutes followed by addition of 20 mM succinic acid as substrate plus 100 mM neutralized hydroxylamine-HCl for 30 minutes at 37 °C. One aliquot of this NIT2 reaction mixture was used for activity assay and another for redox proteomics. The hydroxaminolysis activity assay for NIT2 was performed as previously described.^27^

### Redox proteomics

Free thiols in NIT2 were blocked with 250 mM NEM for 20 minutes and the protein was precipitated with cold acetone overnight at −20 °C. After centrifugation and acetone evaporation, the resulting pellets were solubilized in 6 M guanidinium chloride (GdmCl) and 10 mM tris(2-carboxyethyl)-phosphine (TCEP, freshly added). Samples were further supplemented with 40 mM chloroacetamide, 1 mM CaCl_2_ and 0.01 % ProteaseMAX (Thermo Scientific™ # A40007). Proteins were digested with 1 µg trypsin (sequencing grade, Promega) overnight at 37°C. The digestion was stopped with 1 % trifluoroacetic acid. Purification and elution of peptides was performed as previously described.^46^ After drying, the peptides were resuspended in 10 µl of 1 % acetonitrile, 0.1 % formic acid and stored at −20 °C until MS analysis.

The peptides of each fraction (3 µL) were injected and analyzed by LC-MS/MS using a Q Exactive Plus Orbitrap equipped with an UHPLC Dionex Ultimate 3000 instrument (Thermo Fisher Scientific). The peptides were loaded on an Acclaim™ PepMap™ 100 C18 LC Pre-column (0.1 mm x 20 mm, nanoViper, 5 µm, 100 Å) and separated using emitter columns (15 cm length × 100 μm ID × 360 μm OD × 15 μm orifice tip; MS Wil/CoAnn Technologies) filled with ReproSilPur C18-AQ reverse-phase beads of 3 μm, 100 Å (Dr. Maisch GmbH). HPLC settings: linear gradients of 4 – 25 % acetonitrile (ACN), 0.1 % formic acid (FA) for 35 min followed by 25 – 50 % ACN, 0.1 % FA for 5 min, 50 – 99 % ACN, 0.1% FA for 1 min. The column was washed with 99 % ACN, 0.1% FA for 5 min and then equilibrated with 4 % ACN, 0.1 % FA for 14 min. All flow rates were set as 300 nl·min^−1^. MS data were recorded by DDA. The full MS scan range was 300 to 2000 m/z with a resolution of 70,000 and an AGC of 3E6 with a maximal injection time of 65 ms. The 20 most abundant precursors were selected for MS2. Only charged ions >2 and <8 were selected for MS/MS scans with a resolution of 17,500; isolation window of 2.0 m/z; AGC: 1E5; maximal injection time: 65 ms. MS data were acquired in profile mode.

MaxQuant 2.0.3.0^47^ was used to analyze the MS raw spectra files. The human reference proteome database (UniProt, February 2024, including canonical sequences and isoforms) was used for identification with a false discovery rate (FDR) ≤1%. To quantify the protein abundances, iBAQ values were calculated. To account for protein loading and MS sensitivity variations, the intensities of individual peptides were normalized using the respective iBAQ values of NIT2 from the respective samples.

### Isolation of granulocytes

Neutrophil granulocytes were isolated as previously described.^48^ Zymosan was opsonized with fresh human serum (20 mg/mL) by incubation at 37 °C for 30 minutes. 20.000.000 granulocytes were directly added over to the HUVEC and subsequently activated with opsonized zymosan (3 mg/mL) for 15 minutes at 37 °C.

### CRISPR/Cas9 deletions in HUVEC

Guide RNAs targeting coding sequences for NIT2 and GLS1 were designed using the publicly available CRISPR algorithm (www.benchling.com). gRNAs targeting the regulatory element within the *NIT2* gene were designed using CRISPOR (http://crispor.tefor.net).^49^ gRNAs were cloned into a lentiviral CRISPR/Cas9 v2 (LCV2) plasmid using the “Golden Gate” cloning protocol.^50^ gRNAs were cloned into plasmids containing either puromycin resistance (gift from Feng Zhang, Addgene plasmid #52961; http://n2t.net/addgene:52961; RRID:Addgene_52961) or hygromycin resistance (kindly provided by Frank Schnütgen, Department of Medicine, Hematology/Oncology, University Hospital Frankfurt, Goethe University, Frankfurt, Germany). Lentivirus were produced in Lenti-X 293 T cells (Takara, #632180) using Polyethylenamine, psPAX2 and pVSVG (pMD2.G). pMD2.G was a gift from Didier Trono (Addgene plasmid #12259; http://n2t.net/addgene: 12259; RRID:Addgene_12259). psPAX2 was a gift from Didier Trono (Addgene plasmid #12260; http://n2t.net/addge ne:12260; RRID:Addgene_12260). LentiCRISPRv2-produced virus was transduced for 24 hours in HEK 293 cells or HUVEC (at passage 1) with polybrene transfection reagent (MerckMillipore, #TR-1003-G) and selection was performed with puromycin (1 μg/mL) or hygromycin (100 µg/mL) for 6 days.

#### List of gRNAs including overhangs

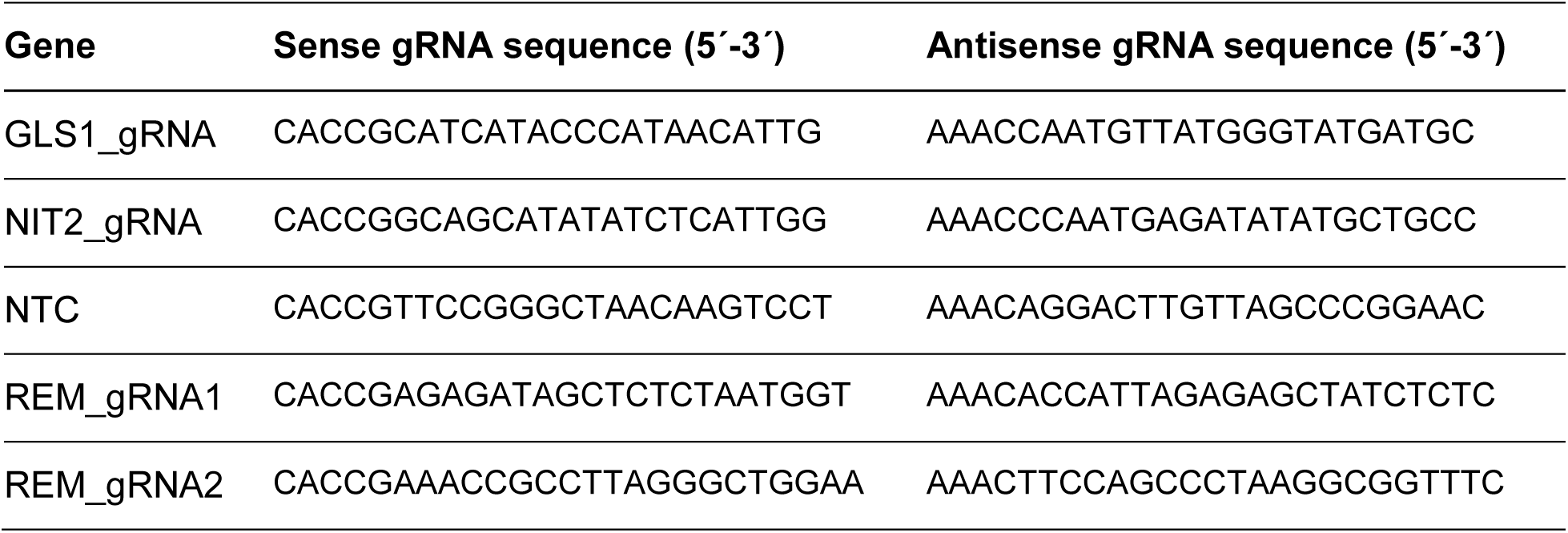

### Western blot analysis

HUVEC were lysed with a triton-based buffer at pH 7.4, with the following concentrations in mmol/L: Tris-HCl (50), NaCl (150), sodium pyrophosphate (10), sodium fluoride (20), Triton X-100 (1%), sodium desoxycholate (0.5%), proteinase inhibitor mix, phenylmethylsulfonyl fluoride (1), orthovanadate (2), okadaic acid (0.00001). Proteins (30 µg) were separated by SDS/PAGE, transferred by Western blot and probed with antibodies as listed below. Western blot analyses were performed with an infrared-based detection system (Odyssey, Licor, BadHomburg, Germany).

#### List of antibodies

| Name | Host | Manufacturer | ID |
| --- | --- | --- | --- |
| β-Actin | Mouse | Sigma-Aldrich,<br>Taufkirchen, Germany | A1978 |
| GLS 1 | Rabbit | Abcam, Cambridge, UK | ab156876 |
| NIT2 | Rabbit | Abcam, Cambridge, UK | ab183074 |
| Anti-His-tag | Rabbit | Bethyl, Texas, USA | A190-114A |

### Immunofluorescence

HUVEC were seeded on Ibidi slides and fixed with 4 % paraformaldehyde that was quenched with glycine (2 %). Next, cells were permeabilized with 0.05 % Triton X-100 in PBS. After blocking with 3 % BSA for 30 minutes, the cells were incubated at 4 °C overnight with a 1:200 dilution of the primary antibody. Cells were washed with 0.3 % Tween 20 in PBS and incubated with a 1:500 dilution of secondary antibody for 30 minutes. The cells were then washed again with 0.3 % Tween 20 and counterstained with DAPI (1:500). Images were captured with a laser confocal microscope LSM800 and analyzed with ZEN lite software.

### Targeted LC-MS/MS analysis for TCA cycle metabolites

#### Metabolite extraction from cells

HUVEC were grown in EGM on a 10 cm dish until confluence. For isotopic labeling experiments, glutamine was replaced with fully labeled glutamine (^13^C_5_, ^15^N_2_). Cells were washed with ice-cold PBS and scratched carefully in PBS. After centrifugation (2,400 x *g*, 4 minutes), the cell pellets were lysed by adding 600 µL of 90 % methanol and two freeze-thaw cycles. After centrifugation, 300 µL of clear supernatant were mixed with internal standards mix containing eight ^13^C-labelled metabolites: α-ketoglutaric acid-1^13^C, citric acid-2-^13^C, glucose-6-phosphate-6-^13^C, D-glucose-1-^13^C, L-glutamic acid-2-^13^C, pyruvate-1-^13^C, succinic acid-2-^13^C and itaconic acid-5-^13^C. Samples were dried and reconstituted in water containing 0.5% formic acid.

#### Metabolite extraction from plasma and urine

Plasma and urine were diluted with 25 µL Trifluorethanol (TFE):H_2_O (2:1) or 25 µL of pure TFE, respectively. After a 10 min incubation, samples received 100 µL of MeOH:EtOH (1:1), followed by 100 µL H_2_O + 10 µL internal standard mix (containing internal Standards homotaurine, citrate-1,5-^13^C_2_ and succinate-1,4-^13^C_2_, all at 0.04 mM). Samples were sonicated, centrifuged and dried under nitrogen flow. Samples were reconstituted in 50 µL 99 % MilliQ water with 1 % MeOH and 0, 2 % Formic acid.

Alternatively, urinary metabolites (human samples) were normalized by the concentration of creatinine, which was determined on diluted urines (1:25, with water) according to a published protocol.^51^ Metabolites were quantified using a 12-point calibration curve created by serial dilution of solutions made with pure analytes dissolved in LC-MS quality water with the following concentration ranges: αKG: 0-500 µM, αKGM: 0-50 µM, and creatinine: 0-500 µM. D_3_-methylmalonic acid (5 µM, for αKG and αKGM) and D_3_-creatinine (25 µM, for creatinine) were used as internal standards. Calibrators, quality controls (urine sample of known creatinine concentration) and study plasma and urine samples were prepared by mixing 20 µL of sample with 20 µL freshly prepared dithiothreitol 0.5 M followed by vortexing and incubation at room temperature for 10 min. 20 µL of internal standard mix were then added and the samples vortexed. Metabolites were extracted by addition of 100 µL of 0.1% formic acid in methanol, followed by vortexing and centrifugation at 10,000 *x g* for 10 min at room temperature. The supernatants were transferred into HPLC vials and 10 µL were injected for LC-MS/MS analysis.

#### TCA-Metabolite analysis

Samples (8 µL) were injected via an Infinity II Bio liquid chromatography system into a 6495C triple quadrupole mass spectrometer (both Agilent Technologies, Waldbronn/Germany). Metabolites were separated on an Acquity HSS T3 C18 column (1.8 µm, 2.1 x 150 mm, Waters) by using the following mobile phase binary solvent system and gradient at a flow-rate of 0.35 mL/min: Mobile A consisted of 100 % water with 0.2 % formic acid. Mobile phase B consisted of 100 % acetonitrile with 0.2 % formic acid. The following 11 minutes gradient program was used: 0 min 1 % B, 0-6 min 1 % B, 6-7 min 80 % B, 7-8 min 80 % B, 8-11 min 1 % B. The column compartment was set to 30 °C. Metabolites were detected with authentic standards and/or via their accurate mass, fragmentation pattern and retention time in polarity switching ionization dynamic MRM AJS-ESI mode, and quantified (where appropriate) via a calibration curve. The gas temperature of the mass spectrometer was set to 240 °C and the gas flow to 19 L/min. The nebulizer was set to 50 psi. The sheath gas flow was set to 11 L/min, with a temperature of 400 °C. The capillary voltages were set at 1000/1000 V with a nozzle voltage of 500/500 V. The voltages of the High-Pressure RF and Low-Pressure RF were set to 100/100 and 70/70 V, respectively. Metabolite peaks were annotated with Skyline-daily (version 22.2.1.278).

For human urine and plasma samples, αKG was determined as previously described^52^ and αKGM was compatible with the chromatographic conditions of the TCA panel. The two fragments of the parent ion (m/z 105,1 Da and m/z 91 Da) were monitored in positive mode with a Sciex 6500+ ESI-tripleQ MS/MS. Signal processing and quantification of metabolites were carried out with Analyst® 1.7.2 software, 2022 AB Sciex.

Metabolites are represented in concentrations when a standard curve was available or their signals were alternatively normalized to that of internal standards and expressed as intensity or peak area.

### Human samples

αKGM and αKG were measured in urine and serum that were collected from a total of 19 males and female individuals (average age of 46) recruited from the employees of the Saarland University Hospital. Only healthy subjects without prevalent cardiovascular or chronic kidney disease, diabetes, and regular medication were included. All participants gave their written consent, and the study was approved by the local institutional review committee (155/13).

### Animal procedure

Nit2-floxed mice were generated by the Laboratory Animal Resource Center, University of Tsukuba as described.^53^ LoxP sites flanking exon 2 of *NIT2* were inserted by means of CRISPR/Cas9. Nit2-knockout mice (Nit2^ko/ko^) were generated by a 597 bp deletion of exon 2 by CRISPR/Cas9. Endothelial cell-specific, tamoxifen inducible knockout mice of NIT2 (ecNit2^−/−^) were generated by crossing Nit2^flox/flox^ mice (Nitrilase-like 2, Nit2^tm1NH^) with Cdh5-CreERT2 (Tg(Cdh5-CreERT2)^1Rha^)^54^ mice (kindly provided by Ralf Adams, Münster, Germany). NIT2 deletion was induced by providing tamoxifen in the diet (400 mg/kg, 10 days) when male animals were at least 8 weeks old. A tamoxifen free “wash out” period of at least 14 days after tamoxifen feeding was adhered to. Control animals (CTL) are defined as Nit2^flox/flox^-Cdh5-CreERT2^0/0^ littermates (i.e. no Cre expression) and were also, treated with tamoxifen.

Systemic inflammation in mice (18 weeks old, C57Bl6/J, 29 ± 3 grams body weight) was induced by a single i.p. lipopolysaccharide injection (LPS from Escherichia coli strain O55:B5, 4 mg/kg, 4h, Sigma-Aldrich, St. Louis, USA).

All animals had free access to chow and water in a specified pathogen-free facility with a 12 hour day/ 12 hour night cycle and all animal experiments were performed in accordance with the German animal protection law and were carried out after approval by the local authorities (Regional council Darmstadt or Northrhine Westfalia, under the approval FU2020 or 81-02.04.2022.A334, respectively). Every mouse received an identification number for each experiment and the experimenter was blind for the genotype or treatment. Animal group sizes differ due to number of littermates. Control and knockout animals were studied in paired fashion per experiment.

### Neonatal retina angiogenesis

One day old pups (ecNit2^−/−^ and their control counterparts) were injected intra-gastricaly with 50 µL of tamoxifen at 1 mg/mL dissolved in sunflower oil for three consecutive days. On day 7 pups (ecNit2^−/−^ or Nit2^ko/ko^ and their respective controls) were euthanized, the eyes were removed and the retina was exposed and divided into four petals by incisions towards the its center. It was fixed with 100% methanol at −20°C overnight. Retinas were washed and treated with 0.1 % Triton X-100, 1 % bovine serum albumin (BSA) and 1 % donkey serum (Sigma, D9663). Next, retinas were stained with isolectin GS-IB4 (1:500, Thermo Fisher, I21411) overnight at 4 °C. Retinas were mounted on microscope slides (Thermo Scientific-Superfrost, 631-9483) using Dako Fluorescence Mounting medium (Agilent Technologies Inc., S3023). Images were acquired with a Zeiss LSM800 laser scanning microscope (Carl Zeiss Microscopy GmbH) under a 20 x objective using the software ZEN (ZEN 3.1, Carl Zeiss Microscopy GmbH). Images were acquired using the tile mode (100 tiles per retina). Quantification of the vascular network was performed using the freely available software Angiotool64 (version 0.6a).

### Aortic ring outgrowth assay

Aortic rings (1 mm) were embedded in a gel mixture of rat-tail collagen I (1.5 mg/mL, #354236, BD) 1x Medium 199 (#M0650, Sigma) and NaHCO_3_ (2.2 mg/mL) at 37 °C for 60 minutes. EBM supplemented with 2.5 % autologous serum was added to the gels. Aortic rings were treated or not with murine VEGF-165 (30 ng/mL, #450-32-10UG, PeproTech) and cultured for 7 days. Rings were fixed with 4 % PFA, treated with 0.5 % Triton X-100 and 1 % BSA. Endothelial cells were staining with an anti-mouse CD31 antibody (1:200, #550274, BD). Images were acquired with a Zeiss LSM800 laser scanning microscope (Carl Zeiss Microscopy GmbH) Images were acquired using the tile- and Z-mode and sprouts were quantified using Angiotool64 (version 0.6a).

### Cell migration

A scratch wound assay was performed with HUVEC in endothelial growth media (EGM) in 96-well plates. Cell migration into the scratched area (Incucyte WoundMaker, wound closure) was monitored by live cell imaging and analyzed using the IncuCyte ZOOM platform.

### Proliferation assay

3,000 HUVEC were seeded out on a 96-well plate in endothelial growth media. Nuclei were stained using the IncuCyte Nuclight Rapid Red Dye according to the manufacturer’s instructions. Proliferation was monitored by live cell imaging using the IncuCyte ZOOM platform.

### Spheroid outgrowth assay

Spheroid outgrowth assays were performed as previously described.^55^ HUVEC spheroids were stimulated for 16 hours with human recombinant VEGF-A (50 ng/mL). Images were generated with the Evos XL Core. The quantitative analysis of sprout number and cumulative length was calculated with the AxioVision software.

### RNA-sequencing

RNA and library preparation integrity were verified with LabChip Gx Touch 24 (Perkin Elmer). Sequencing was performed on NextSeq2000 instrument (Illumina) with 1 x 72 bp single end setup. The resulting raw reads were assessed for quality, adapter content and duplication rates with FastQC (RRID:SCR_014583).^56^

Sequencing reads were aligned against the hg38 genome assembly using STAR (v2.7.10, RRID:SCR_004463), with the parameter – quantMode set to “GeneCounts”. Differential gene expression analysis was performed using DESeq2 (v1.32.0; RRID:SCR_015687) in R (v4.1.1; R Project for Statistical Computing (RRID:SCR_001905).^57,58^ Differentially expressed genes were taken as those with a false discovery rate-adjusted p-value of less than 0.05. Gene set enrichment analysis was performed using the gprofiler2 (v0.2.3, RRID:SCR_018190) package for R, using the WikiPathways (RRID:SCR_002134) source. F1 scores were calculated using the resulting precision and recall values, as standard. Visualizations of the data and results were generated using the R package ggplot2 (RRID:SCR_014601) in R.

### β-Galactosidase assay

HUVEC were cultured in endothelial basal media (EBM) with 2 % FCS for 16 hours. Cells were fixed and treated following manufacture’s instructions in the Senescence β-Galactosidase Staining Kit (#9860, Cell Signaling Technology). Senescence is expressed by the percentage of β-galactosidase positive cells counted in at least 3 randomly chosen images per each of the three batches of HUVEC.

### Statistics

Data are represented as mean ± standard error of the mean. Calculations were performed with Prism 9.2.0. The latter was also used to test for normal distribution and similarity of variance. In the case of multiple testing, a Bonferroni correction was applied. For multiple group comparisons, analysis of variance followed by post hoc testing was performed. Individual statistics of dependent samples were performed by paired t-test, of unpaired samples by unpaired t-test, and, if not normally distributed, by the Mann-Whitney *U* test as indicated. P values of <0.05 were considered significant. Unless otherwise indicated, *n* indicates the number of individual experiments. Statistics for RNAseq, MS and metabolomics were carried out as described in the specific sections.

---

**Suppl. Figure 1:**
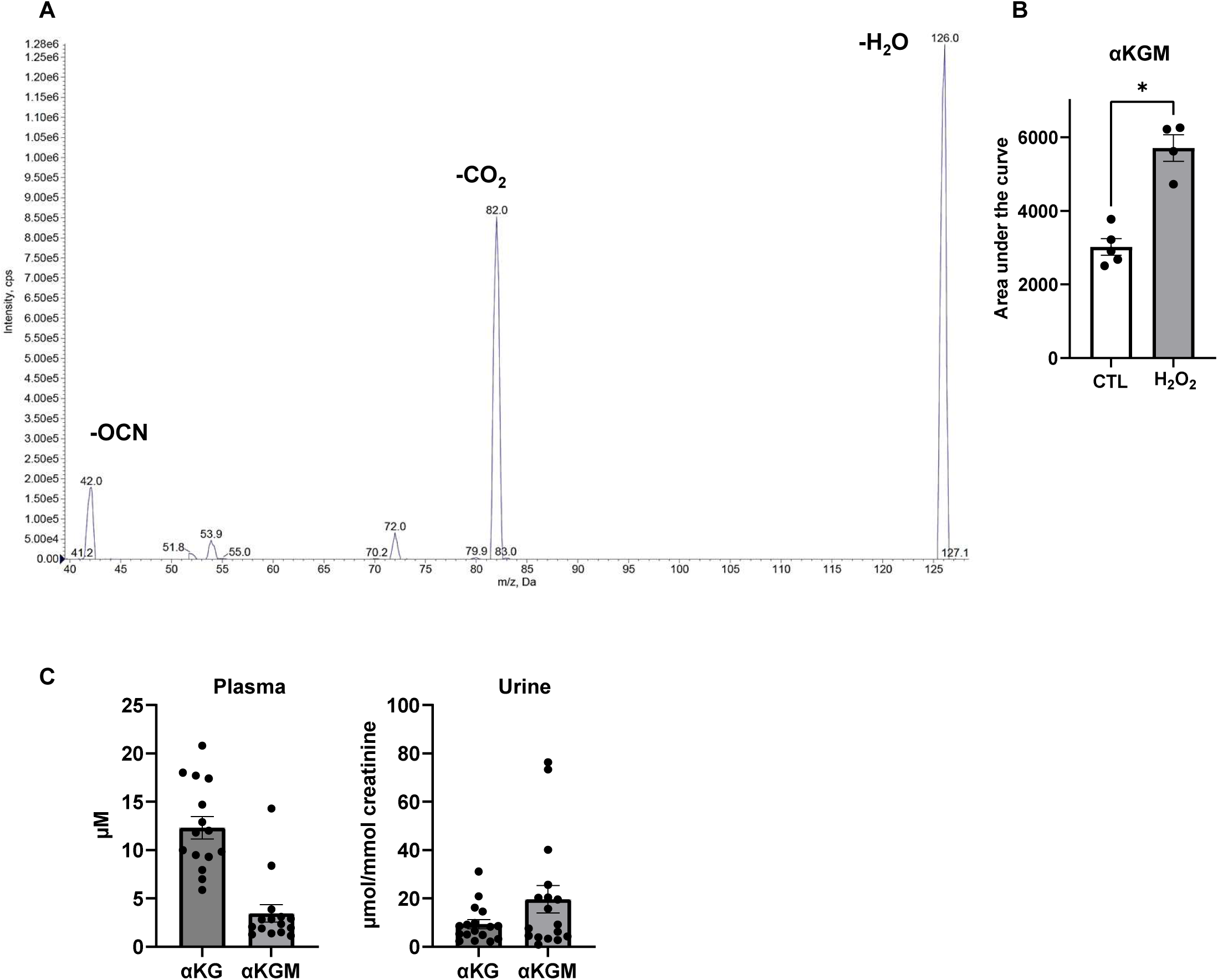
Targeted LC-MS/MS of αKGM. A: The mass spectrum of αKGM in its lactam form as detected in negative ionization mode. The spectrum shows three major fragmentation peaks at 126.0 Da (M-1-H_2_O; −18 Da), 82.0 Da (M-1-CO_2_-H_2_O; −62 Da) and 42.0 Da (a CNO-fragment). B: Targeted LC-MS/MS measurements for intracellular αKGM in HUVEC with or without exposure to H_2_O_2_ (300 µM, 15 minutes). C: αKG and αKGM concentrations (using calibration curves of known concentrations of each compound) in plasma and urine (normalized to creatinine) of healthy individuals.

**Suppl. Figure 2:**
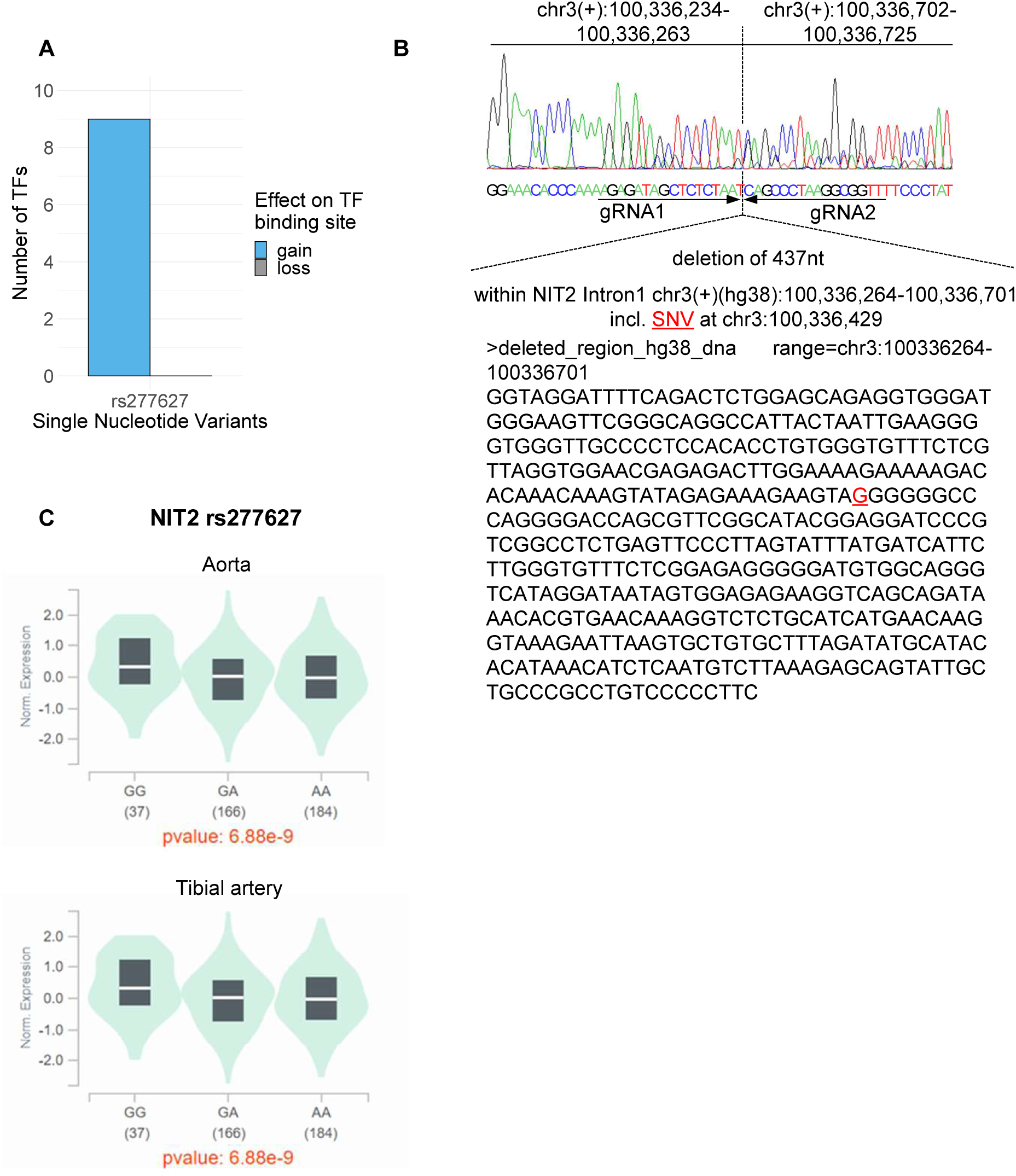
Regulation of *NIT2* expression by the SNV rs277627. A: rs277627 is located in a regulatory element (REM, analyzed with EpiRegioDB) with binding sites for the transcription factors ZNF320 VEZF1 KLF16 GLIS2 MAZ ZNF740 ZNF148 ZNF467 GLIS3 that are expressed in HUVEC and have a gain of function and likely repress transcription. B: A two gRNA approach was employed to delete a 437 bp region where the SNV rs277627 (chr3:100336428-100336429) in NIT2 (intron 1) is located. gRNA1 targets the chr3:100,336,234-100,336,263 region whereas gRNA2 the chr3:100,336,702-100,336,725. A successful deletion was obtained between 100,336,264-100,336,701 as shown by sanger sequencing. The SNV G is labeled in red. C: Data from GTex for the expression of wild type (GG) rs277627 and its variants GA and AA on NIT2 expression in human aorta and tibial artery.

**Suppl. Figure 3:**
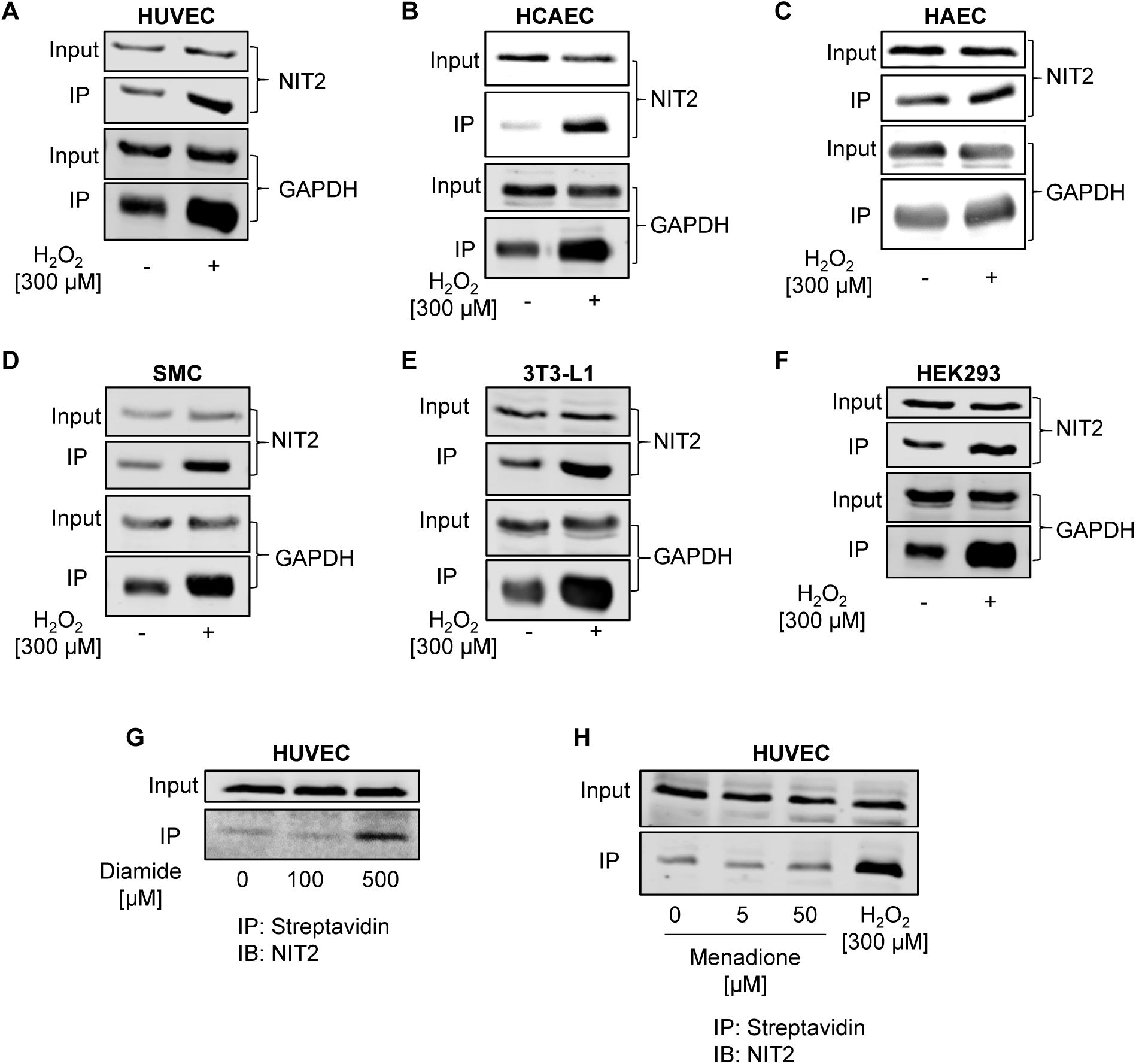
Cysteine oxidation in NIT2. Biotinylated iodoacetamide (BIAM) switch assay in different cell types exposed or not to 300 µM H_2_O_2_, 15 minutes. (A) HUVEC, (B) HCAEC, (C) HAEC, (D) SMC, (E) mouse fibroblasts 3T3-L1, (F) HEK 293. BIAM switch assay in HUVEC exposed to diamide (G) or menadione (H). HUVEC: human umbilical vein endothelial cells HCAEC: human coronary artery endothelial cells HAEC: human aortic endothelial cells SMC: smooth muscle cells IP: immunoprecipitation IB: immunoblotting GAPDH: Glyceraldehyde-3-Phosphate Dehydrogenase

**Suppl. Figure 4:**
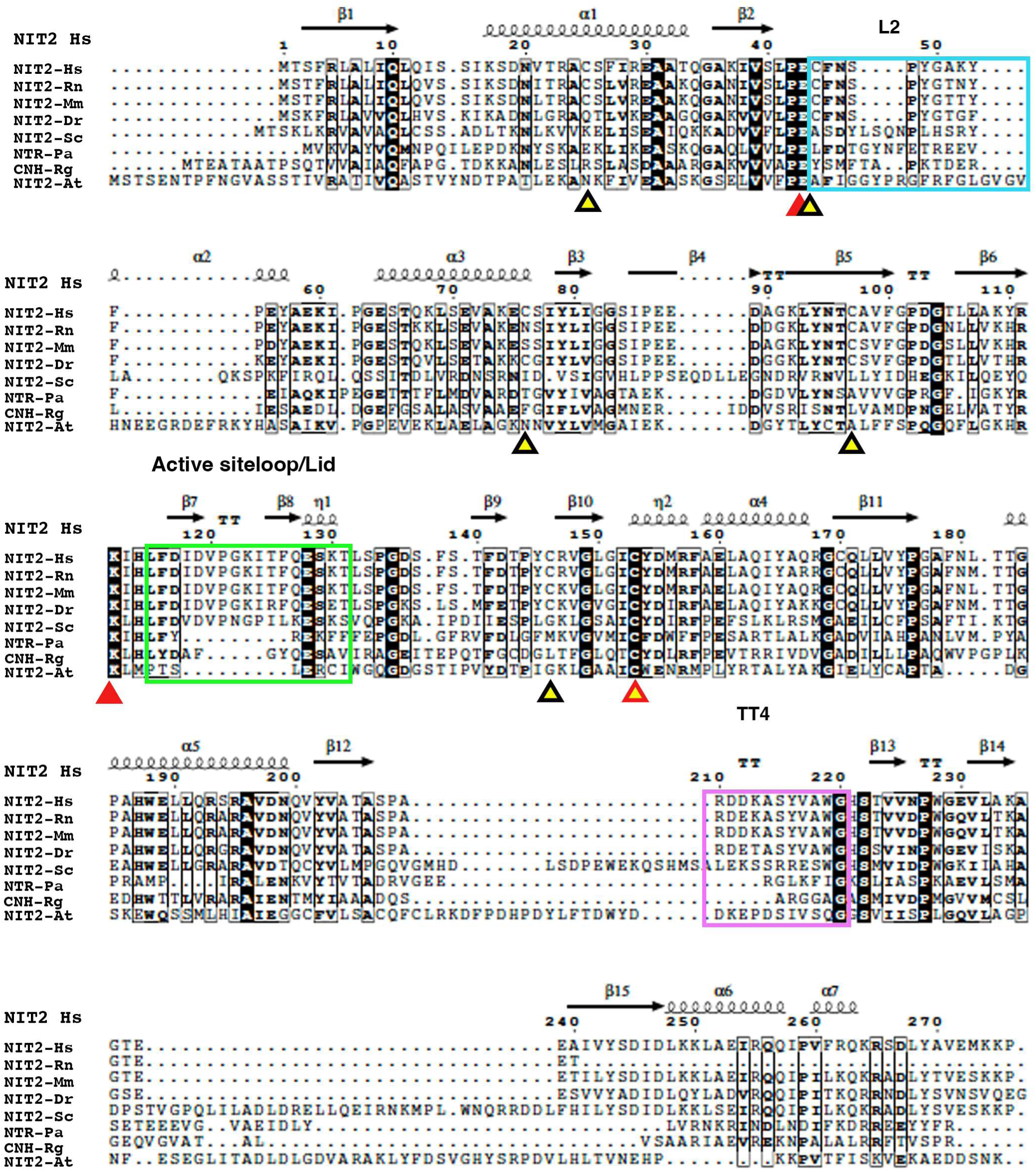
Animal NIT2 contains more cysteine residues than that of other organisms or plants. Multiple sequence alignment and structural features of human NIT2 predicted by AlphaFold2. Abreviations: Hs, *Homo sapiens* (Q9NQR4); Rn, *Rattus norvegicus* (Q497B0); Mm, *Mouse musculus* (Q9JHW2); Dr, *Danio rerio* (Q4VBV9); Sc, *Saccharomyces cerevisiae* (P47016); Pa, *Pyrococcus abyssi* (Q9UYV8); Rg, *Rhodococcus qingshengii* (A0AA46MND0); At, *Arabidopsis thaliana* (P32962). The human cysteine residues are indicated with yellow triangles, the catalytic residues as red triangles, and the loops involved in the formation of the substrate channel and active site are colored in blue, green, and pink squares.

**Suppl. Figure 5:**
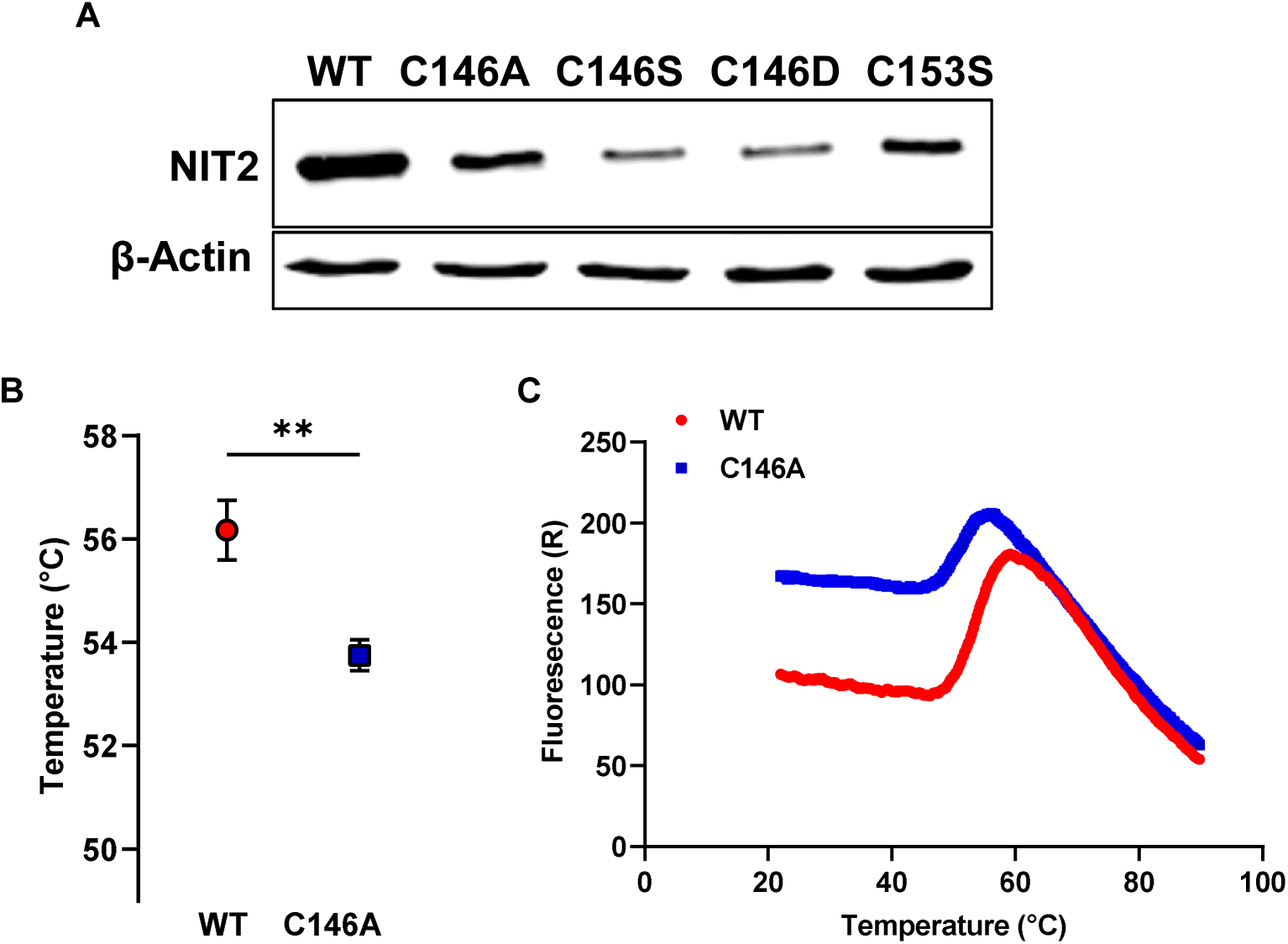
Expression and stability of NIT2 C146 mutants. A: Western blot analysis (using an anti-His-tag antibody) for the His-tagged mutants of NIT2 C146 expressed in HEK 293 cells. B: Aggregation points of NIT2 and NIT2 C146A as determined by thermal shift assay. n=3; ** p<0.01, Welchs’ correction. C: Melting curve of purified NIT2 and NIT2 C146A protein.

**Suppl. Figure 6:**
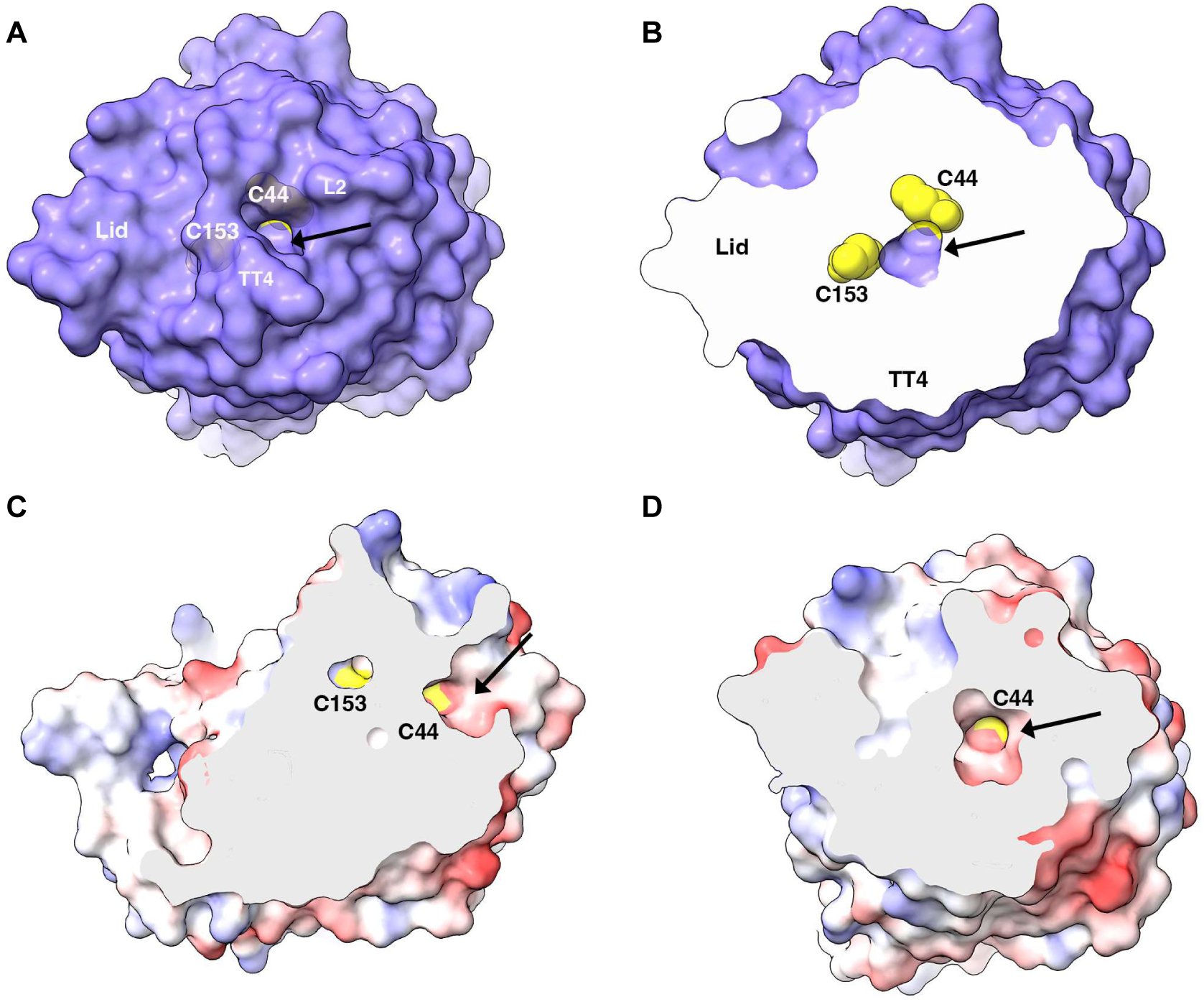
Cysteine 44 forms the substrate channel and is prone to oxidation. The AlfaFold2 structure prediction of NIT2 is depicted as the solvent-exposed surface in purple, and the substrate channel entry is shown from the top view (A) and the cut view to the bottom (B). The surface charge representation (red negative, white hydrophobic and blue positive) is shown on the side cut view (C) and top cut (D). Cys153 and Cys44 depicted as yellow spheres. The black arrow indicates the channel entry.

**Suppl. Figure 7:**
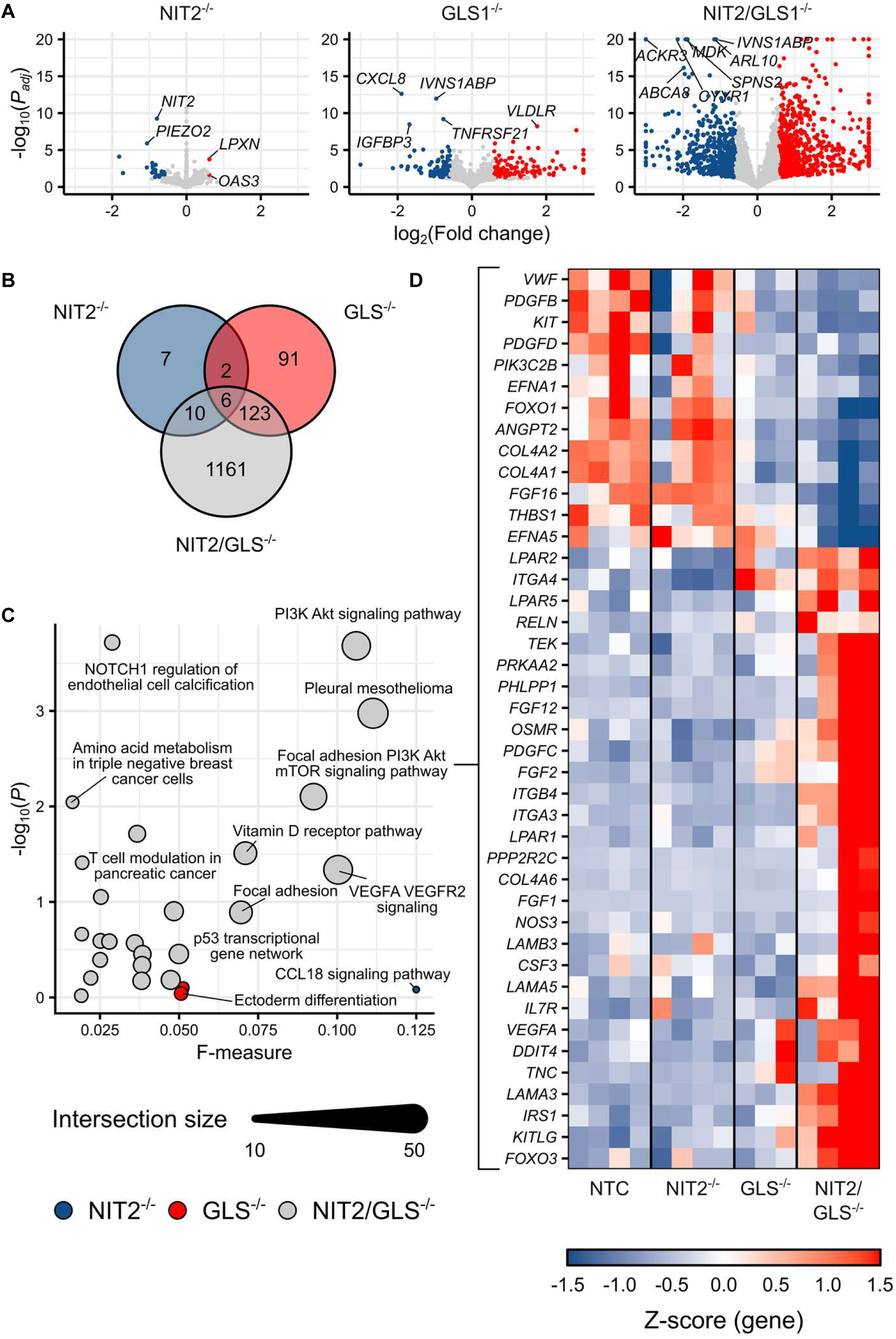
NIT2 and GLS1 sinergistically control gene expression (RNAseq) in HUVEC. A: Volcano plots for each knockout as indicated. B: Number of differentially expressed genes. C: Annotation pathway. D: Heat map of top differentially expressed genes in HUVEC NTC, NIT2^−/−^, GLS1^−/−^ and NIT2/GLS1^−/−^.

**Suppl. Figure 8:**
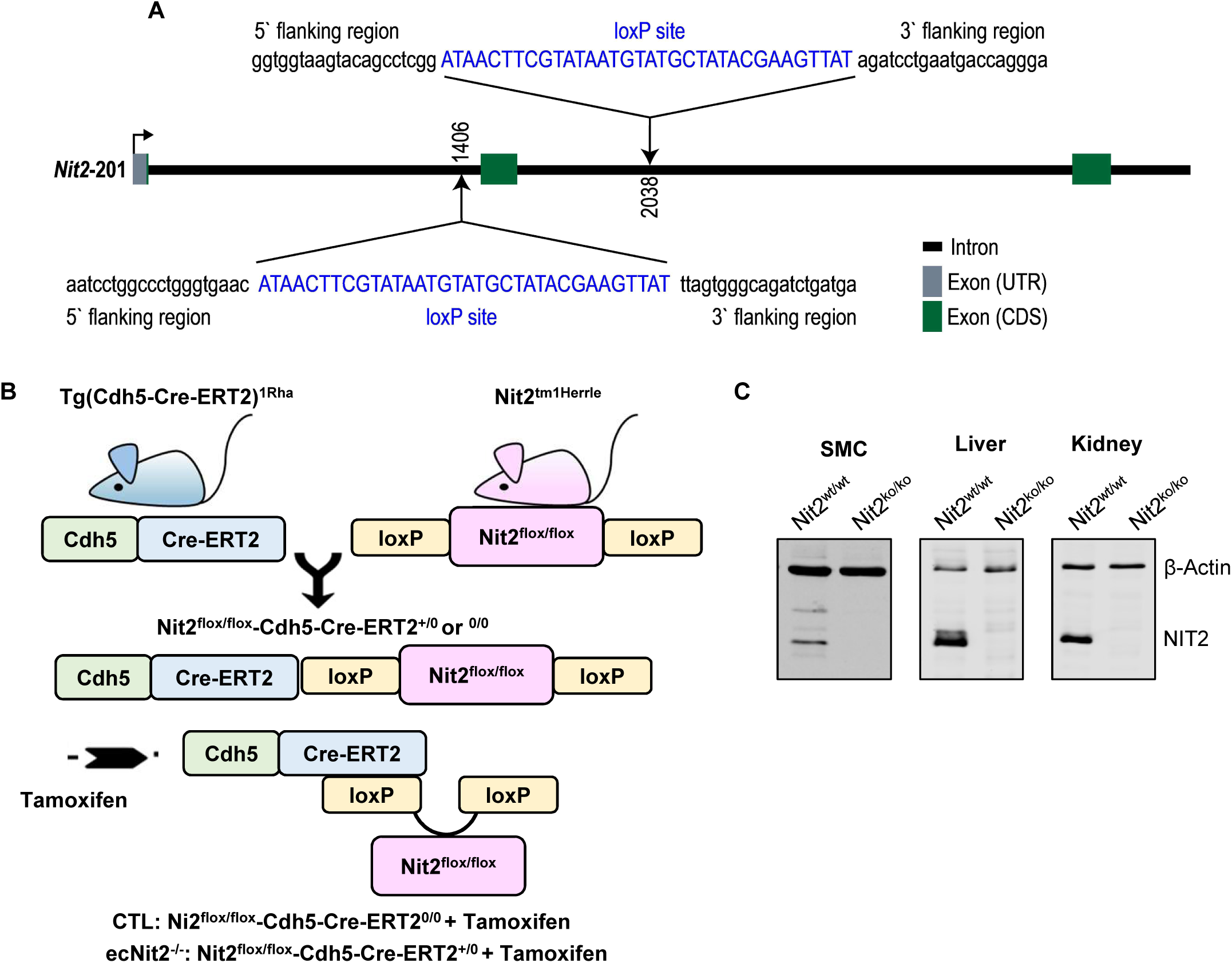
Generation of Nit2 knockout mice. A: LoxP sites flanking exon 2 of the *NIT2* gene were inserted by CRISPR/cas9. B: Nit2^flox/flox^ mice were crossed with Tg(Cdh5-Cre-ERT2)^1Rha^ to generate endothelial-specific, tamoxifen-inducible knockout mice of Nit2. C: Validation of knockout efficiency by Western blot analysis for Nit2 in global, constitutive knockout mice of Nit2 that were generated by CRISPR/cas9 deletion of ∼500bp. Nit2^wt/wt^ and Nit2^ko/ko^ are littermates in a heterozygous breeding of Nit2^wt/ko^ with Nit2^wt/ko^ mice. SMC: aortic smooth muscle cells.

## Notes

### Competing Interest Statement

The authors have declared no competing interest.

